# The single-cell spatial landscape of stage III colorectal cancers

**DOI:** 10.1101/2024.11.07.622577

**Authors:** Andrew Su, HoJoon Lee, Minh Tran, Richard D. Cruz, Anuja Sathe, Xiangqi Bai, Ignacio Wichmann, Lance Pflieger, Bryce Moulton, Tyler Barker, Derrick Haslem, David Jones, Lincoln Nadauld, Quan Nguyen, Hanlee P. Ji, Terence Rhodes

**Affiliations:** Division of Oncology, Department of Medicine, Stanford University School of Medicine, Stanford, CA, 94305, United States; Institute for Molecular Bioscience, The University of Queensland, QLD 4072, Australia; Intermountain Healthcare, Saint George, UT, 84770, United States; Division of Obstetrics and Gynecology, Department of Obstetrics, Escuela de Medicina, Pontificia Universidad Católica de Chile, Santiago, 8331150, Chile

## Abstract

We conducted a spatial analysis using imaging mass cytometry applied to stage III colorectal adenocarcinomas. This study used multiplexed markers to distinguish individual cells and their spatial organization from 52 colorectal cancers. We determined the landscape features of cellular spatial features in the CRC tumor microenvironment. This spatial single-cell analysis identified 10 unique cell phenotypes in the tumor microenvironment that included stromal and immune cells with a subset which had a proliferative phenotype. These special features included spatial neighborhood interactions between single cells as well as different tissue niches, especially the tumor infiltrating lymphocyte regions. We applied a robust statistical analysis to identify significant correlations of cell features with phenotypes such as microsatellite instability or recurrence. We determined that microsatellite stable (MSS) colorectal cancers had an increased risk of recurrence if they had the following features: 1) a low level of stromal tumor-infiltrating lymphocytes, and 2) low interactions between CD4+ T cells and stromal cells. Our results point to the utility of spatial single-cell interaction analysis in defining novel features of the tumor immune microenvironments and providing useful clinical cell-related spatial biomarkers.

## INTRODUCTION

Colorectal cancer (**CRC**) is a leading cause of cancer-related deaths. An important contributor to CRC development, maintenance and metastasis is the tumor microenvironment (**TME**). This cellular milieu contains neighboring tumor epithelial cells, normal epithelium, fibroblasts, endothelial cells, immune cells and many others. The characterization of the TME is critical for understanding tumor biology as well as determining specific features with clinical implications. Representing a subset of the TME, the tumor immune microenvironment (**TIME**) has specific cellular characteristics that are important predictors for the response to treatments such as immunotherapy.

Immunohistochemistry (**IHC**) and immunofluorescence (**IF**) are commonly used assays applied to tissue sections. IHC and IF detect specific protein cancer markers such as immunotherapy targets like PD-1 and PD-L1. These immune checkpoint markers provide spatial cellular information about the tumor microenvironment and its immune cell components. However, methods such as IHC and IF have major issues that include significant variation in their staining patterns and limits in the number of protein markers^1,2^. With the introduction of new spatial imaging methods, one can characterize the TIME with single-cell precision and much greater multiplexing capacity^3^. For example, imaging mass cytometry (**IMC**) provides the spatial detection from eight to 120 proteins from a given tissue sample and resolves the expression from individual cells^3^. Spatial imaging methods have identified new breast cancer subtypes and the spatial properties of the TIME, some of which may be linked to clinical outcomes and immune response^4,5^. Spatial analysis with single cell resolution can infer cell-cell interactions as a proxy for TIME states and activities in the tumor - these features may be useful to measure disease progression and response to specific therapies^6^.

The prognostic value of tumor-infiltrating lymphocytes (**TILs**) has been extensively studied^7,8^. Currently, pathologists’ visual inspection is the most used approach for identifying and roughly estimating the number of TILs. However, the manual characterization of TILs is a challenging task, prone to high level of interobserver variability and not scalable for large number of samples^9^. A lack of reproducibility limits the use of these conventional means for measuring the presence of TILs^10,11^. Moreover, histopathology assessment of TILs is qualitative and does not account for spatial features such as proximity to other cell types. Approaches such as IMC provide a more objective, quantitative approach for spatially assessing TILs. Overall, spatial assays for TIL measurement will provide significant insights into the CRC TIME.

For this study, we focused on a set of stage III CRCs – these patients have CRC with local lymph node involvement. Currently, clinical stage is the most important factor in terms of ascertaining prognosis. The 5-year overall survival rate for CRC varies by stage at diagnosis, ranging from 90% for early-stage (I-II), to 70% for locally advanced stage (III) and to less than 15% for metastatic (IV)^12^. Even after surgical resection, patients with limited stage (I-III) CRCs have an increased risk of metastatic recurrence. Adjuvant chemotherapy for stage III CRC decreases the risk but the recurrence of metastasis is still high with approximately 20% of patients have new sites of CRC after treatment^13^. Given the poor prognosis of CRC metastasis, there have been multiple studies to identify specific molecular and cellular markers identifying the stage III patients at highest risk^14–16^.

Herein, we evaluated spatial data from primary colorectal cancers, characterizing cell-cell interactions and describing the spatial patterns of cell neighborhoods. We developed some novel approaches for evaluating spatial data, leveraging the single cell resolution of spatial features. We applied this approach to this set of stage III CRCs, determined their spatial TIME features and characterized cellular interactions among the various cells. Finally, we explored the potential of integrating IMC-based, spatially-derived TIME features were associated with metastatic recurrence.

## RESULTS

### A computational framework for analyzing single-cell spatial features in CRC cancer

We performed IMC assays on a set of stage III CRC samples to achieve high-resolution, single-cell spatial profiling (**Fig. 1a**). The IMC multiplexed panel utilized 16 antibodies that are markers for the following: 1) tumor epithelial cell markers such as E-cadherin and TP53, 2) immune cell markers such as CD3, CD4+, and CD8a, 3) stromal cell markers such as SM-actin and collagen, and 4) nuclear markers such as the Histone H3 protein (**Supplementary Table 1**).

**Figure 1.**
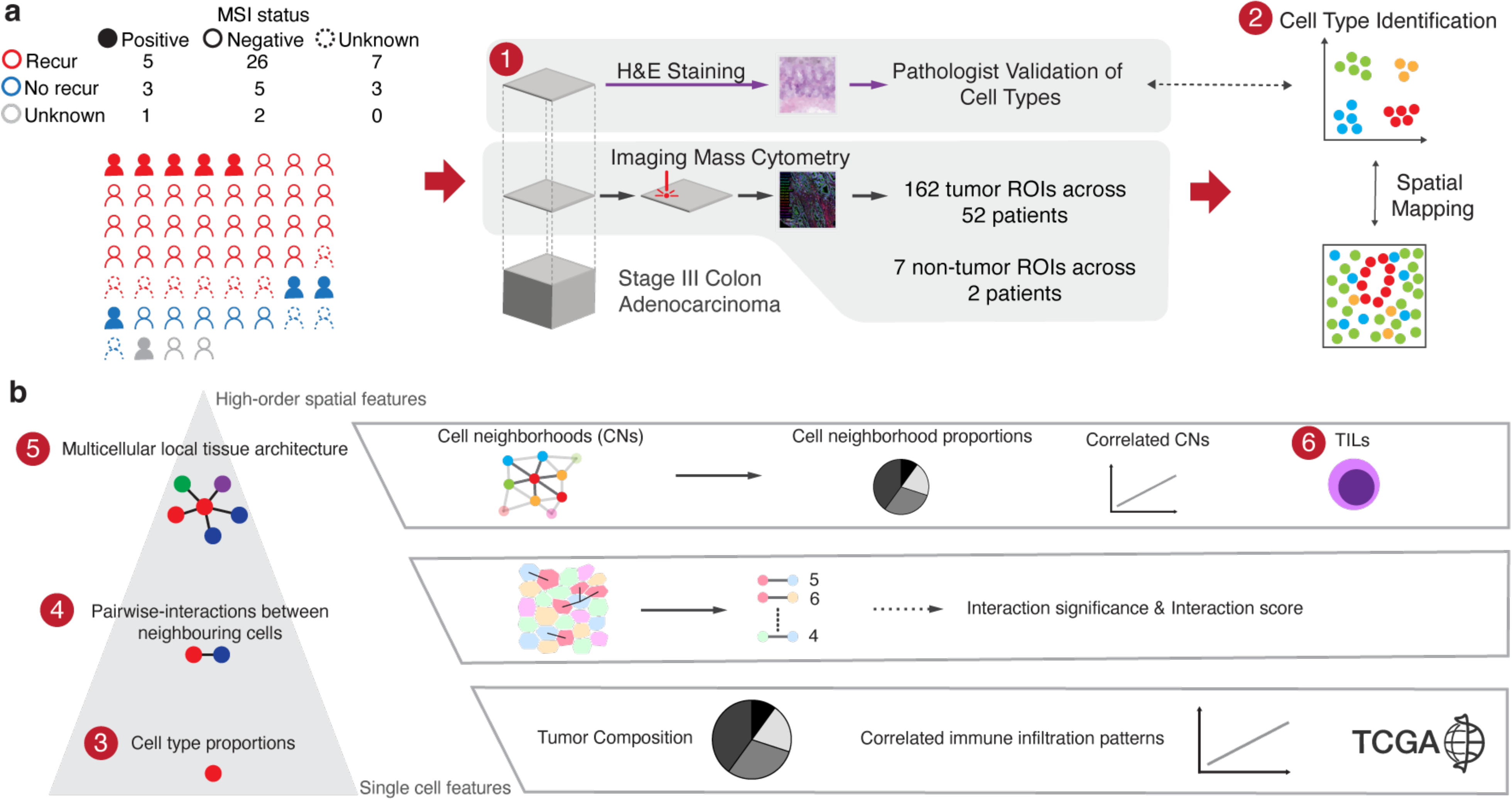
Overview of the computational framework for single-cell spatial analysis. **a)** Cohort overview and data generation pipeline, b) Analytical workflow: from single-cell features to higher-order spatial characteristics.

To analyze this single-cell level spatial data from IMC, we developed an analytical framework (**Fig. 1b**) with three tiers: 1) delineating cell type proportions, 2) investigating pairwise interactions among neighboring cells, and 3) characterizing the multicellular local tissue architecture within cellular neighborhoods (**Methods**).

We used spatial IMC data from the 52 CRCs covering 162 tissue regions and seven non-tumor tissue regions from normal colon. We analyzed these tissue regions, with image dimensions ranging from 141 µm × 500 µm to 1121 µm × 1309 µm. The sample image count per each sample, ranged from 1 to 8, allowing us to cover more area from a given tissue (**Supplementary Fig. 1a**). Other molecular characteristics such as tumor purity and number of cells per tissue regions are summarized in **Supplementary Figure 1**.

The first level involved single cell analysis - we examined the cell types and determined their proportions (**Fig. 1b**). The results included quantitative characteristics of immune cell infiltration abundance across different cell types for a given cancer tissue region. The second level involved cellular organization - we investigated the spatial relationships among the various cell types. We measured the distances between cells and co-occurrence of different cell types using built-in functions in two programs, Histocat^17^ and Squidpy^18^. Afterwards, we conducted statistical comparisons to determine of the enrichment of cell type pairs and their colocalization. Based on the comparison of the observed colocalization to the null distribution, we identified changes in interactions between neighboring cells across samples and infer potential interactions contributing to the immune response. The third level involved spatial cellular organization in a tissue - we evaluated thousands of local regions, each composed of up to the 10 cell types^19^. We refer to these regions as cell neighborhoods (**CNs**) where all cells are located within 40 µm radius (see Methods). Thus, we identified the spatial arrangement and potential interactions of various cell types within the localized tissue regions. These cell neighborhoods helped us define and pinpoint TILs in the tumor microenvironment.

Subsequently, we determined if there were correlations among spatial cellular features and various clinical metrics including microsatellite instability (**MSI**) status and disease recurrence, typically meaning the identification of a metastasis after surgery and adjuvant chemotherapy (**Supplementary Table 2**). MSI is an indicator of loss of DNA mismatch repair – tumors with these features respond to immune checkpoint inhibitors. All CRC samples had genomic data available including exome sequencing^20^. We determined the MSI status using the program MSIsensor^21^ on the exome data (**Supplementary Fig. 2**). For a subset of the samples, we had the results from separate IHC assays for DNA mismatch repair proteins to determine microsatellite status. There was complete concordance between the two MSI metrics.

### Single cell identification of cell types in stage III CRC

The cohort included 52 stage III CRC with additional clinical information such as MSI and recurrence status (**Supplementary Table 2**). After surgical resection of their CRCs, majority of patients had received chemotherapy as adjuvant treatment. For each CRC undergoing IMC analysis, we identified individual cell types that included tumor, immune, and stromal cells. After preprocessing and cell segmentation with 16 different antibodies, we had a total of 903,125 cells for all samples. Subsequently, we quantified the expression intensity of each protein marker from the cells and identified clusters through unsupervised Leiden clustering (**Supplementary Fig. 3a**). The clusters showed ten specific cell types as shown visually using Uniform Manifold Approximation and Projection (**UMAP**) (**Fig. 2a, Supplementary Fig. 3b**)^22^.

**Figure 2.**
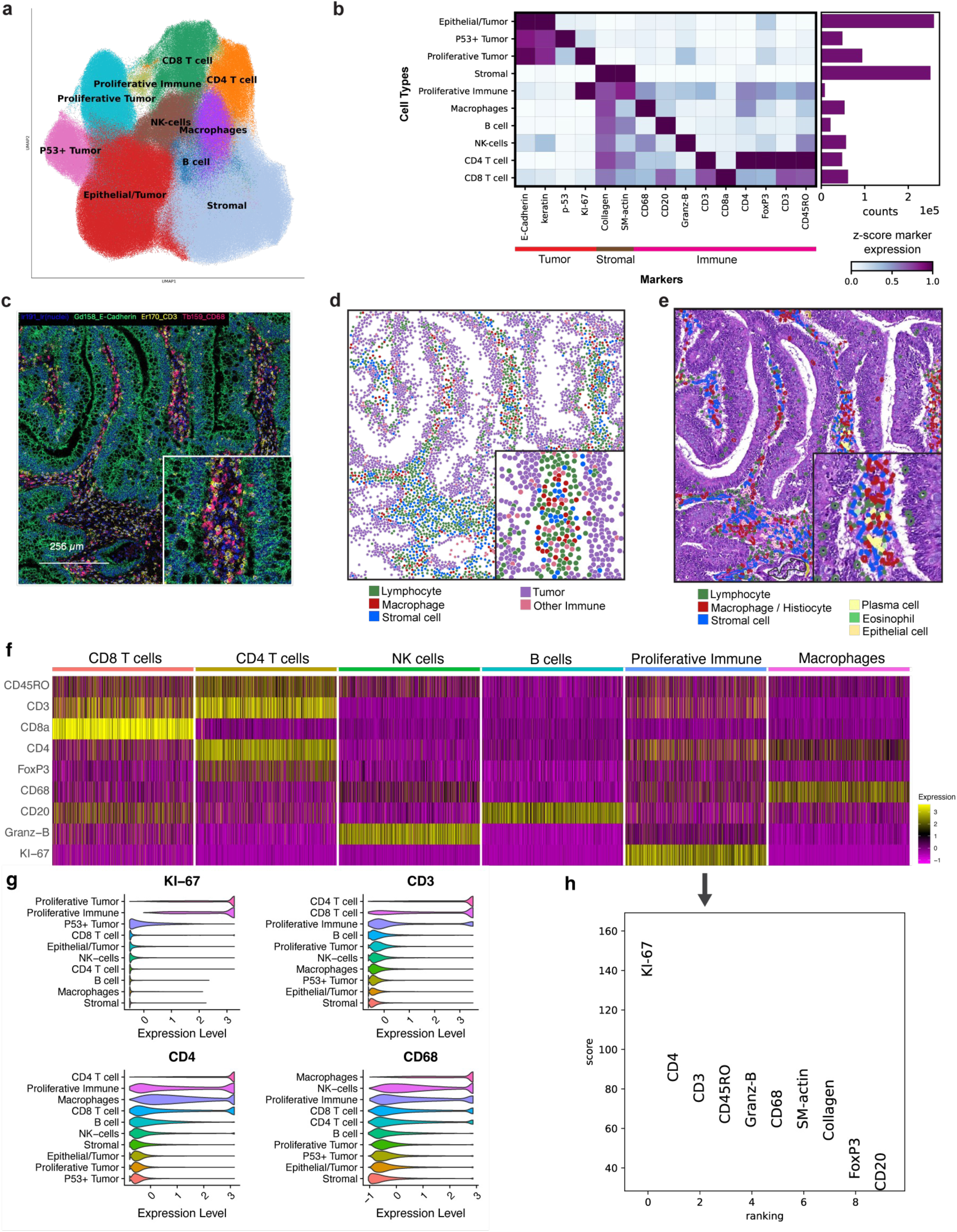
Determining single-cell identity. **a)** The UMAP plot displaying 10 cell types identified via unsupervised Leiden clustering, b) heatmap of lineage marker expression for each cell type, accompanied by a bar plot showing the total cell count for each type across all samples, c) example of an IMC Hyperion image, d) corresponding cell type annotation image, e) corresponding H&E image with pathologist annotations, f) heatmap of immune cell makers for various immune cell types, g) the expression level of four markers across the 10 cell types, f) relative expression of markers in proliferative immune cell type compared to the other cell types.

From this set of samples, we identified 10 different major cell types with several different subcategories (**Fig. 2b**). The non-immune included epithelial tumor cells, TP53+ tumor cells, proliferative tumor (Ki-67+) and stromal cells. For the classification of immune cell types, we used established markers^23^ (**Fig 2f, Supplementary Table 1**). The immune types included proliferative immune, macrophages, B cells, NK cells, CD4+ T cells and CD8+ T cells. The most prevalent cell types were tumor epithelial and stromal cells which were most abundant in CRC tumor regions^24,25^. To corroborate our results, a pathologist conducted a visual inspection of cells embedded within the tissues with histopathologic annotation. This independent review confirmed the cell type classifications as we determined from the IMC data (**Fig. 2c, d, e**).

Macrophages were the most abundant immune cell type, a finding that is supported from prior reports^26^. We identified proliferative immune phenotypes, representing a rare population of immune cells (<1% of total cells) distinguished by their high Ki-67 expression (indicating their proliferative state) along with expression of other immune markers (**Fig. 2g**). Cells undergoing proliferation were identified by high KI-67 expression – this class of proliferating immune cells are increasingly being reported among various studies^27–29^. Additional immune markers indicated a prominent prevalence of CD4+ T cells, accompanied by T cells and macrophages, within the population of proliferative immune cells (**Fig. 2h**).

### Different cell proportions across the CRCs and their TMEs

We determined the distribution and abundance of the 10 unique cell types for the CRCs (**Fig. 3a**). This analysis included determining the abundance of the immune cell types as a fraction of total immune cells (**Fig. 3b**). For comparison as an independent data set, we evaluated the cellular content of 77 stage III CRC samples of the Cancer Genome Atlas (**TCGA**). For these CRCs, Luca *et al*. estimated the abundance of various cell types from bulk RNA-seq using the CIBERSORTx approach^25^. Between these two sets of stage III CRCs, we observed the same ranking of abundance for the different immune cell types (**Supplementary Fig. 4**). The high level of concordance between these two independent sets of CRCs validated our cell type identification using spatial analysis. We also estimated tumor purity (**Supplementary Fig. 1b,c**), employing it as a covariate in our linear models in downstream analyses. This step allowed us to control for any sampling bias of tissue regions in the ensuing analyses.

**Figure 3.**
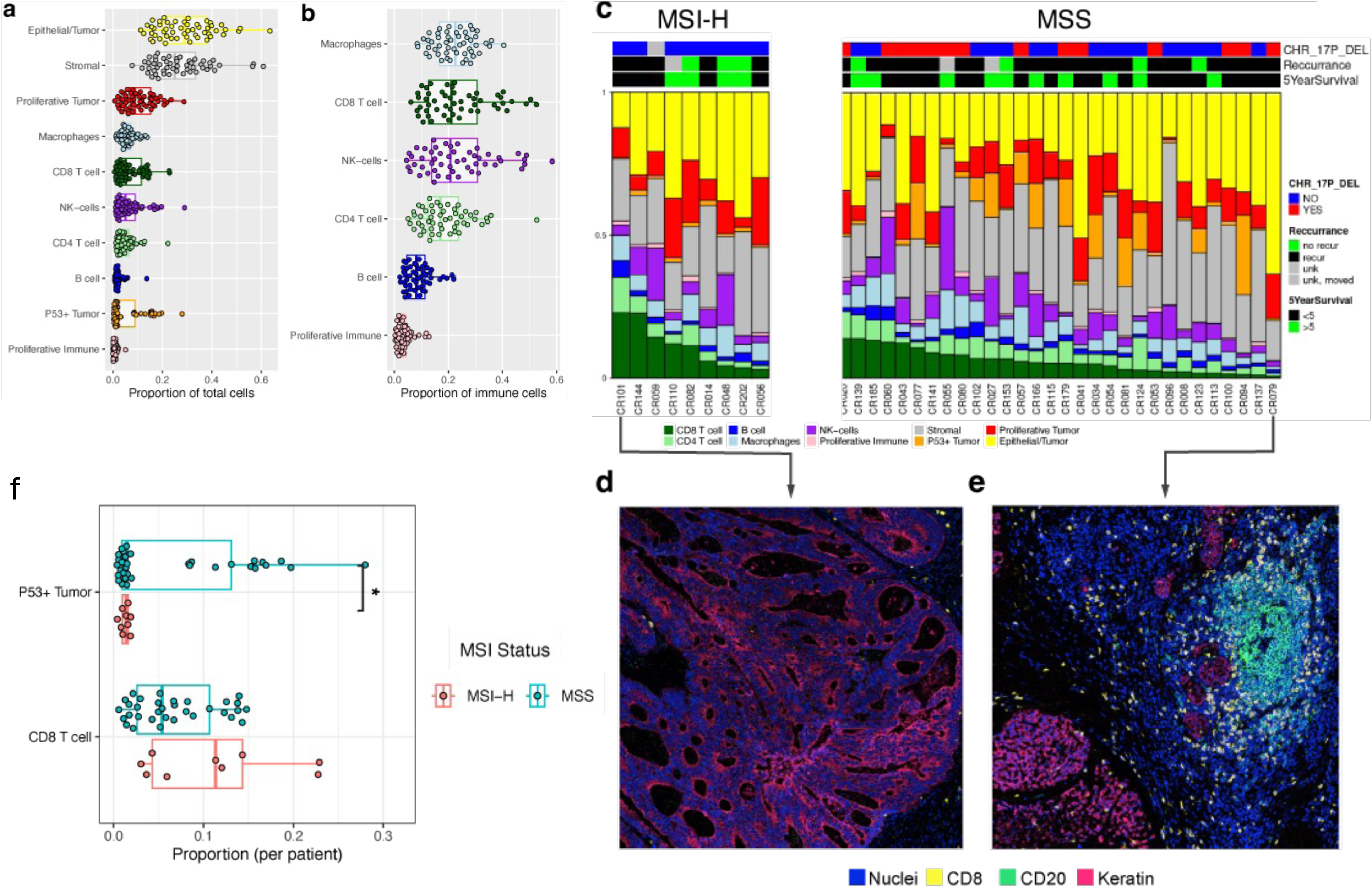
Cell type composition across all samples. **a)** The box plots showing the distribution of all cell type proportions, b) immune cell type proportions across all samples (n = 52), c) stacked bar plot illustrating cell type proportions in MSI (n = 9) and MSS (n = 33) samples, ordered by CD8+ T cell proportion, d) representative IMC images of tissue region from a patient with the highest CD8+ T cell proportion, e) a patient with the lowest CD8+ T cell proportion, f) box plots comparing the proportion of TP53+ tumor cells and CD8+ T cells between MSI and MSS tumors (*p < 0.05), g) cell type proportion correlation correlograms for patients with tumor recurrence (MSI: n = 5, MSS: n=26), showing only significant correlations (Spearman’s correlation, adjusted p ≤ 0.05). Color scale bar indicates the strength and direction of correlations across tissue regions.

### Cellular composition among CRCs with microsatellite instability

Approximately 10-15% of stage III CRCs have an important molecular phenotype related to loss of DNA mismatch repair (**MMR**)^30^. Loss of this repair mechanism leads to a hypermutable state called microsatellite instability (**MSI**)^30^. Mutations occur at a much higher rate in DNA sequences called microsatellite tracts. High levels of MSI are also referred to as ‘MSI-H’ whereas microsatellite stable is called ‘MSS’. In general, MSI-H CRCs have a better prognosis compared to MSS CRCs^31,32^. MSI-H CRCs have an associated higher levels of TILs^33^. In addition, MSI in CRCs is a clinical predictor for immune checkpoint inhibitors.

Among the CRCs with MSI status, nine had MSI-H and thirty-three were MSS. Subsequently, we compared the cellular composition between the MSI-H and MSS tumors. We arranged samples by the abundance of CD8+ T cells as shown in **Figure 3c**. Examination of the IMC images showed extensive heterogeneity in cell type composition. For example, the MSI-H CRC (CR101) had a high proportion of these cells (**Fig. 3d**), whereas a MSS CRC (CR079) had a much low proportion of CD8+ T cells (**Fig. 3e**). Citing another example, MSS CRCs like CR060 and CR055 had an increased number of NK cells.

There were some specific trends in cellular composition between MSI-H and MSS samples (**Supplementary Fig. 5a,b**). Prior studies have shown that tumors with *TP53* mutations have a higher level of TP53 protein staining^34–36^. Moreover, MSI tumors have a significantly lower frequency of *TP53* mutations (∼7%) compared to MSS tumors (∼71%)^37^. Among our tumors, MSI-H CRCs had a significantly lower proportion of TP53^+^ tumor cells than MSS CRCs (adjusted p value=0.038, **Figure 3f**). In fact, fewer than 5% of cells in MSI-H tumors were TP53^+^ tumor cells (**Supplementary Fig. 5b**). We also corroborated this relationship in terms of tumor cells with TP53 protein expression using genomic data from our prior publication^20^. We also observed that CRCs with *TP53* missense mutations had significantly more TP53^+^ tumor cells compared to CRCs that were wildtype *TP53* (**Supplementary Fig. 5c**).

Among the other MSI-H CRC features there was a higher proportion of CD8+ T cells compared to MSS samples (p value = 0.047, adjusted p value = 0.234) (**Fig. 3f**). This result is consistent with other studies^38–40^. For MSI-H tumors, the increase in CD8+ T cells is attributed to their having a higher level of immunogenicity that reflects a greater number of cancer mutations that generate neoantigens compared to MSS CRCs^41,42^. From the TCGA CRC (**COAD**) cohort, there were higher numbers of CD8+ T cells in MSI-H tumors versus MSS tumors among stage III CRCs (**Supplementary Fig. 6b**).

Next, we evaluated correlations between the proportions of pairs of distinct cell types in MSS tumors. For instance, we investigated whether CRCs with a low proportion of CD4+ T cells also had a low proportion of CD8+ T cells. Subsequently, we determined if there were correlations based on CRCs from patients with recurrence versus those without recurrence (**Supplementary Fig. 7**). We observed a significant positive correlation between the proportions of CD4+ and CD8+ T cells among recurred patients’ tumors (adjusted p-value < 0.05), where both CD4+ and CD8+ T cell proportions were low. This result aligns with observations from previous reports^43,44^. In contrast among patients without recurrence, there was no correlation between these two cell types.

Additionally, there were several other significant correlations between different cell types observed only in MSS CRCs from patients with recurrence (**Supplementary Fig. 7a**). Notably, the proportion of proliferative tumors was significantly higher in patients with right-sided colon cancer compared to those with left-sided colon cancer (adjusted p-value = 0.045). It is worth mentioning that more right-sided CRCs had a higher frequency *TP53* missense mutations compared to left-sided ones, and all nine MSI patients had right-sided colon cancer.

### Cell interactions in the CRC TME

We leveraged spatial information to identify pairwise interactions between adjacent neighboring cells. We defined interacting cells as being 6 µm or less apart between two adjacent cell borders (e.g., membrane) (**Supplementary Fig. 8a**). We employed two methods for identifying significant pairwise interactions: 1) a permutation-based neighborhood approach described in Histocat to identify global patterns of interactions (**Fig. 4a**); 2) a method that we developed which provides a quantitative interaction score which we used to assess spatial relationships with specific clinical metrics (**Fig. 4b**).

**Figure 4.**
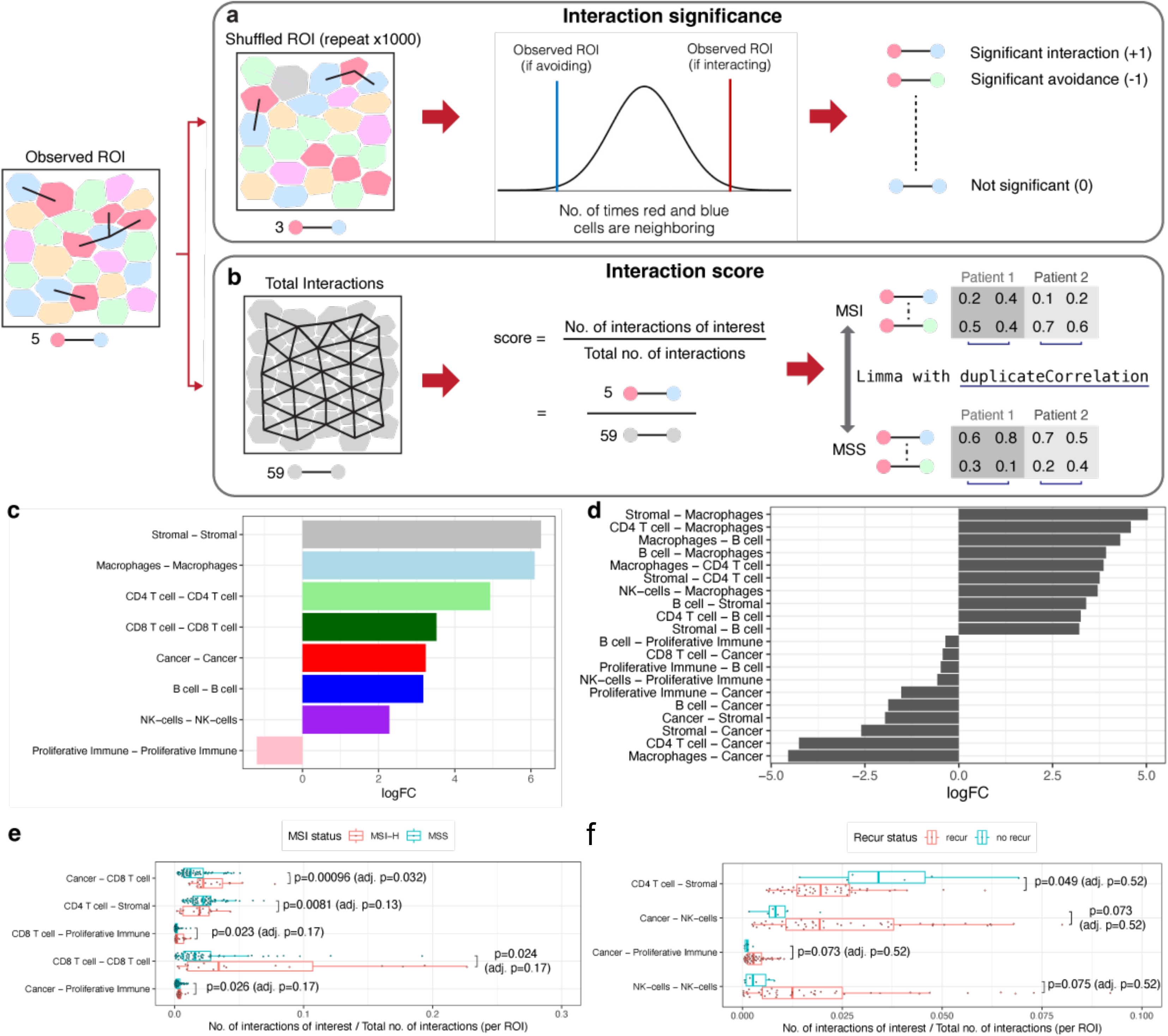
Cell-Cell interactions in the tumor microenvironment. **a)** An overview of Histocat’s random permutation approach for assessing significant interactions or avoidance between cell type pairs at the tissue region level, b) quantitative interaction scores for identifying significant pairwise cell interactions between two conditions, c) homotypic interactions shown as the logarithmic fold-change (**LogFC**) of interacting versus avoiding tissue regions, d) heterotypic interactions highlighting the top 10 most engaged and avoided pairs, e) boxplots comparing interaction scores between MSI-H and MSS samples, displaying the 5 lowest adjusted p-values, e) boxplot of interaction scores between two cell types in MSS samples, comparing cases with and without recurrence.

Histocat identifies cell type to cell type interactions for specific regions-of-interest from a given spatial image^17^. Briefly, this algorithm compares the number of observed interactions against an expected distribution, computed through random shuffling of cell types. This information is used to determine which cell type pairs preferentially interact versus non-interacting pairs that are separated spatially (**Fig. 4a, Supplementary Fig. 8**). We summarized these results across all tumors as the log fold-change (**logFC**) between the number of interacting and avoiding regions-of-interest for each cell type pair (**Supplementary Fig. 9**).

We determined the homotypic and heterotypic interactions. The homotypic interactions refer to the same cell types being adjacent to each other. The heterotypic interactions refer to where different cell types are adjacent. We observed that homotypic interactions were prevalent across all cell types, with the notable exception of the proliferative immune cell class (**Fig. 4c**). This unique pattern can be attributed to proliferative immune cells engaging with each other in MSI-H tumors but lacking this type of interactions in MSS tumors (**Supplementary Fig. 10a**). Among the heterotypic interactions, we observed a trend where immune cells had a higher degree of close spatial interactions with stromal cells while being more distant from cancer cells (**Fig. 4d**). This interaction pattern was particularly notable among macrophages, CD4+ T cells, and B cells.

The Histocat method also provides a ‘directionality’ metric for each pairwise interaction. This metric is defined based on a pair of different cells where the first type is surrounded by the second type. Our results revealed a spatial arrangement where cancer cells were frequently surrounded by immune cells, including lymphocytes (**Supplementary Fig. 9c,d**).

### Quantitative interaction scores between different neighboring cells

Histocat determines the significant pairwise interactions within individual tissue regions. However, it provides only a categorical output with the following labels: significant interaction, significant avoidance, or non-significance. Categorical outputs are not as amenable to determine the statistically significant pairwise interactions across multiple tissue regions.

Additionally, there is a lack of spatial analysis methods that provide statistical comparisons while treating regions-of-interest correctly as replicates of a patient’s tumor and not as independent samples. To overcome these intrinsic limitations of Histocat, we developed a quantitative interaction score, defined as the proportion of interactions between a specific pair of cell types versus the interactions among all types of neighboring cells (**Fig. 4b**). Like other proportion-based comparisons, this approach enables: 1) rigorous statistical testing for significantly different interactions between conditions; 2) evaluating different regions-of-interest from the same sample.

With this method, we determined if there were interactions among the different cell types in the CRCs (**Fig. 4e,f)**. Among the MSI-H CRCs, CD8+ T cells interacted with proliferative immune cells (p value = 0.023, adjusted p value = 0.17). This proliferative subclass of immune cells was predominantly composed of CD4+ T cells. Among the MSS CRCs, we observed a trend where CD4+ T cells were more likely to interact with stromal cells like fibroblasts (p value = 0.0081, adjusted p value = 0.13).

### Cell neighborhood categories in the immune microenvironment

We quantified patterns in the local cellular architecture defined as cell neighborhoods (**CNs**). For the initial step in characterizing CNs, we identified each individual cell’s ten closest neighbors to define the local cellular microenvironment. For nearly all cells, the 10 nearest neighbors were located within a 40 μm radius. This metric is also an indicator of direct cell-cell interactions within the target cell’s proximal microenvironment^15^. With K-means clustering we identified and quantified the different CN classes for each central cell and the composition of its ten nearest neighbors (**Methods**). A total of eight major CN classes were identified: 1) P53+ tumor; 2) stromal; 3) bulk tumor; 4) immune-enriched stromal; 5) CD8+ T cell enriched; 6) proliferative tumor; 7) NK-cell enriched; 8) mixed immune. For example, a CD8+ T cell could be classified as belonging to a ‘bulk tumor’ CN if epithelial tumor cells constitute the most of its neighboring cells. Similarly, a CD4+ T cell is assigned to a ‘stromal’ CN if stromal cells represent the largest proportion of its ten nearest neighbors (**Fig. 5a**).

**Figure 5.**
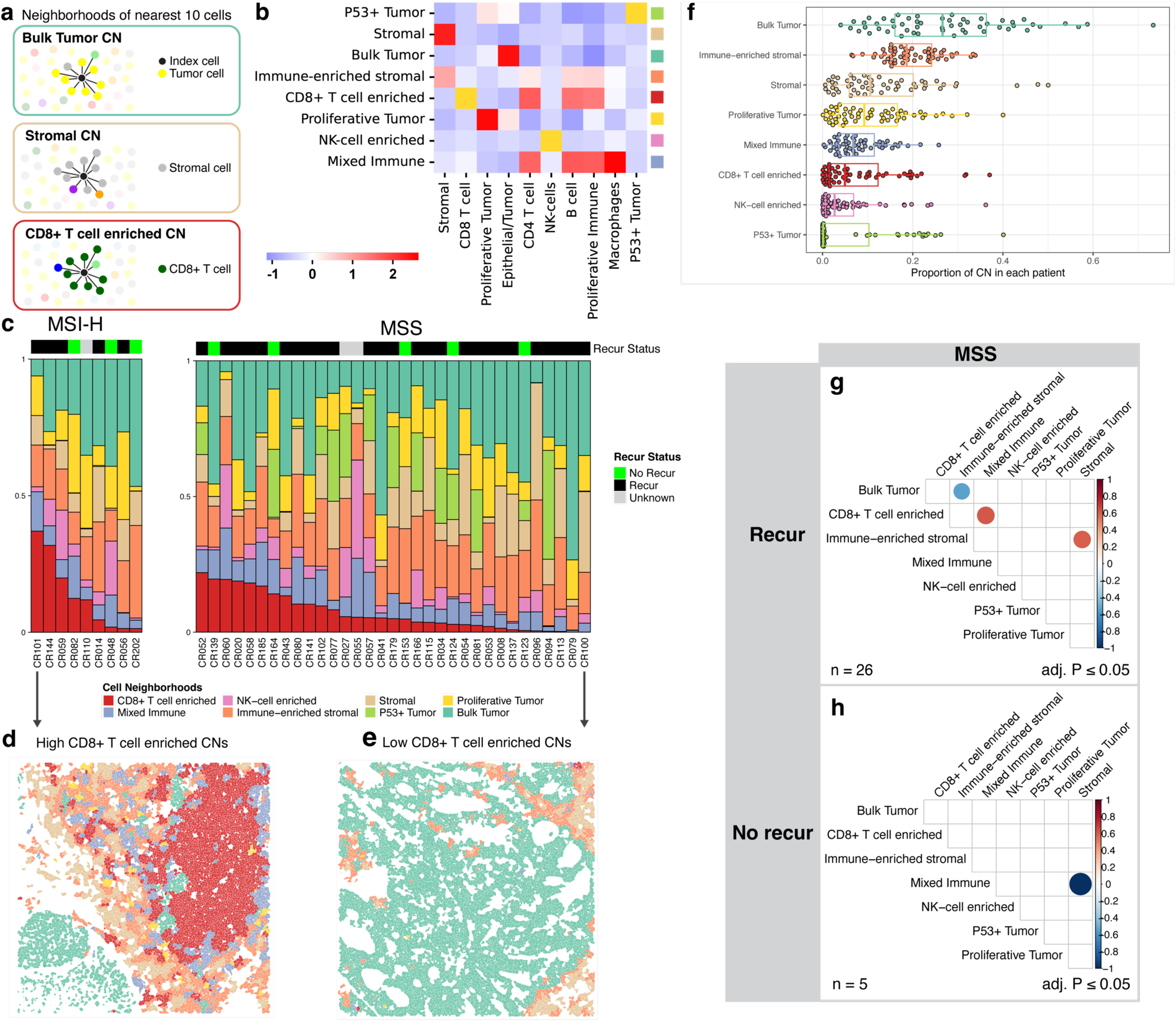
Cell neighborhoods (CN) in Stage III colorectal cancer. **a)** definition of cell neighborhoods with three examples, b) K-means clustering identifies 8 distinct CNs, with a heatmap showing the cell type composition for each cluster (x-axis: cell types; y-axis: CNs), c) Stacked bar plot of CN composition in each patient’s tumor, categorized by MSI status and ordered by CD8+ T cell-enriched CN proportion, d) Example tissue region from a patient with the most abundant CD8+ T cell-enriched CNs, e) from a patient with the least abundant CD8+ T cell-enriched CNs, f) Overall abundance of each CN across all samples, g and h) correlation correlograms between different CN proportions among MSS patients with and without recurrence, only showing significant relationships (adjusted p ≤ 0.05). The scale bar indicates correlation strength across tissue regions.

Each single cell was assigned to the one of eight CNs based on their neighbors (**Fig. 5b**). As expected, the CD8+ T cell-enriched CNs were predominantly composed of CD8+ T cells (>40%), accompanied by other immune cell types such as CD4+ T cells (**Supplementary Fig. 11a**). Similarly, the proliferative tumor CNs were primarily composed of dividing tumor cells (>50%), that were adjacent to non-proliferating epithelial/tumor cells (**Supplementary Fig. 11a**). These patterns were consistently identified across all CRCs.

We determined the representation of CNs such as the abundance of CD8+ T cell-enriched CNs (**Fig. 5c**). To illustrate the range of CN spatial patterns, we show contrasting tissue regions from two different CRCs. The two tumors show the spectrum of CN abundance scale for CD8+ T cell-enrichment. **Figure 5d** shows a tissue region from CRC101, an MSI+ tumor which exhibited the highest abundance of CD8+ T cell-enriched CNs in our cohort. **Figure 5e** depicts a tissue region from CRC100, a MSS tumor which displayed the lowest abundance of CD8+ T cell-enriched CNs. For this CRC, the bulk tumor CN was the dominant type. The distribution of CN proportions across all CRCs is shown in **Figure 5f**. The bulk tumor CNs were the dominant major feature, as expected. Immune-related neighborhoods, including CD8+ T cell-enriched CNs, were frequently present as well.

### Paired cell neighborhood proportions and recurrence

We evaluated the possible correlations between the different cell neighborhood pairs (**Supplementary Fig. 12**). For example, we compared the proportion of the CD8+ T cell CN versus the stromal CN. Afterwards, we determined if the CN correlations were different between CRCs patients with recurrence versus no recurrence. We conducted this analysis across all CRCs and then examined the MSS subset, not having MSI, for associations in recurrence. There were too few MSI samples to determine any CN correlations with recurrence.

However, we successfully assessed the proportion of lymphocytes such as CD4+ and CD8+ T cells within tumor CNs between MSI-H CRCs and MSS CRCs. MSI-H CRCs had a trend towards higher levels of CD8+ T cells, CD4+ T cells, and B cells within tumor CNs compared to MSS CRCs (**Supplementary Fig. 11b**). The proportion of proliferative immune cells within tumor CNs was significantly higher in MSI-H tumors compared to MSS tumors (adjusted p-value = 0.0438, **Supplementary Fig. 11b**). This finding was corroborated by interaction score analyses, which revealed that proliferative immune cells had stronger interactions with cancer cells in MSI-H tumors compared to MSS tumors (**Fig. 4e**).

We examined if specific CN types were associated with recurrence among MSS CRCs. For patients with recurrence, we observed a significant negative correlation between the abundance of immune-enriched stromal CNs and the bulk tumor CNs (adjusted p-value < 0.05) (**Fig. 5h**).

For patients without recurrence, we identified a significant negative correlation between mixed immune CNs and stromal CNs (adjusted p-value < 0.05). Finally, we examined the abundance of proliferative tumor CNs, as a single variable. This CN did not show any differences between the recurrence versus non-recurrence CRCs. This result suggests that CN pairs may be more informative when looking for specific clinical associations.

### Evaluating intratumoral and stromal tumor infiltrating lymphocytes

We used the CN method for characterizing the spatial properties of TILs. Within the local tumor microenvironment, lymphocytes can be spatially associated and interact with either cancer or stromal cells. To quantify these TIL interactions, we defined two types (**Fig. 6a**): 1) intra-tumoral TILs where lymphocytes were located within tumor-dominated CNs (e.g., bulk tumor, proliferative tumor, and TP53+ tumor CNs); 2) stromal TILs where the lymphocytes were located within stromal CNs as characterized by a high proportion of fibroblasts. **Figure 6b** illustrates the cellular composition and TIL distribution across patients. The top panel shows the proportion of lymphocytes, tumor CN cells, and stromal CN cells. The middle and bottom panels depict the percentage of intra-tumoral TILs within tumor CNs and stroma TILs within stromal CNs, respectively, for each patient. The proportion of intra-tumoral TILs among tumor CNs ranges from 0.007 to 0.294 while the proportion of stroma tils among tumor CNs ranges from 0.049 to 0.304.

**Figure 6.**
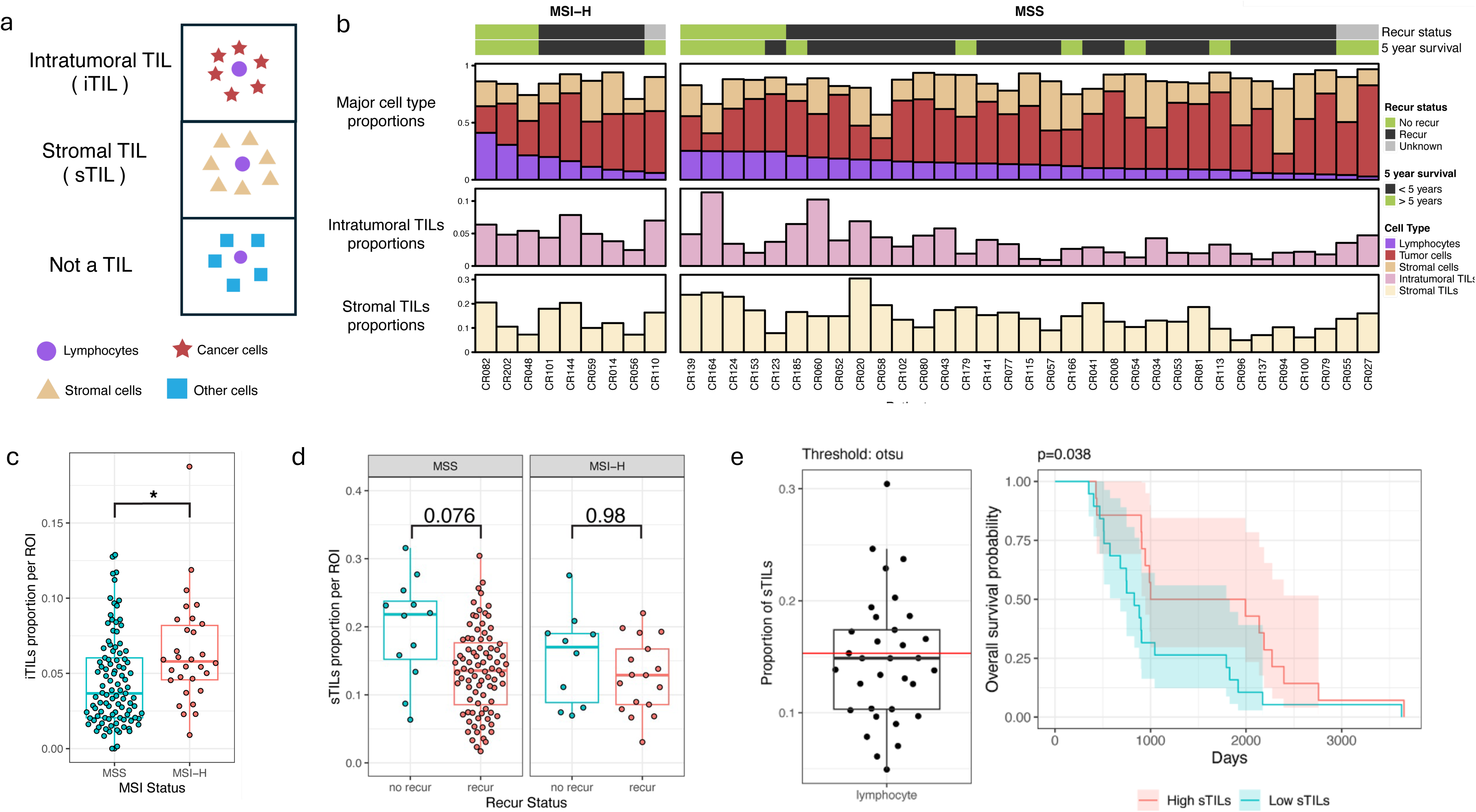
Spatial definition and clinical relevance of TILs in CRC Stage III. **a)** definition of lymphocytes as two types of TILs: intra-tumoral TILs (intra-tumoral TILs) in tumor CNs and stromal TILs (stroma tils) in stromal CNs, b) major cell type proportions including intra-tumoral TILs and stroma TILs across all samples with MSI status (n=42), c) intra-tumoral TIL proportion differences per tissue region between MSI and MSS samples, d) stromal TIL proportion differences per tissue region between recure and no recur among MSS or MSI samples, e) classification of samples into high and low stroma tils using Otsu thresholding, with corresponding overall survival curves for all MSS samples (n = 33).

### Correlation of intratumoral and stromal tumor infiltrating lymphocytes with CRC features

There was a significantly higher proportion of intra-tumoral TILs in MSI-H versus MSS tumors (p value = 0.036, **Fig. 6c**). The addition of spatial CN features was important in recognizing this difference between MSI-H versus MSS CRCs. For example, if we used simple TIL quantitation, e.g., the number of lymphocytes within a given region without spatial metrics, there was no significant difference observed between the MSI-H and MSS CRCs (**Supplementary Fig. 12b**). This result highlights the importance of considering spatial context when characterizing immune cell distributions in the TIME.

To validate these findings, we obtained an independent spatial image data set from 35 advanced stage CRCs^15^. This study used a different approach for protein spatial analysis, co-detection-by-indexing (**CODEX**). Conducted by Schürch *et al.*, they relied on a 56-protein marker CODEX assay applied to 27 primary CRCs from Stage III patients and 7 CRCs from Stage IV patients^15^. This study had 140 CRC tissue regions. The cohort included four MSI+ cases among 27 Stage III samples. Since this study did not report results related to MSI status, we applied our spatial analysis methods to the data, incorporating metrics such as intra-tumoral TILs, as previously described.

From this independent CODEX CRC dataset, we observed similar trends despite the differences in the number of markers (16 vs. 56). First, the rank order of immune cell proportions was consistent, with macrophages being the most abundant cell type, followed by CD4+ T cells, and so on. Second, Schürch et al. also found that the most frequent pairwise cell-cell contacts were homotypic. While their cell-cell contact analysis did not detect interactions between CD4+ T cells and stromal cells, they reported an enrichment of CD4+ T cells in CN-3 (immune-infiltrated stroma) among CLR patients with longer survival compared to DII patients with shorter survival. This finding aligns with our results, where our interaction score revealed stronger interactions between CD4+ T cells and stromal cells in patients without recurrence compared to those with recurrence (**Fig. 4f**). To identify TILs per our criteria, we classified lymphocytes located in CN-2 (bulk tumor) as intra-tumoral TILs. We observed a subtle trend of higher intratumoral lymphocyte presence in MSI-H tumors based on our iTIL definition, whereas overall lymphocyte proportions were lower among MSI tumors in this cohort. However, these observations were not statistically significant due to the small sample size of MSI cases (n=4, **Supplementary Fig. 13**).

Next, we analyzed the stromal TIL distribution in relation to recurrence status and MSI. Patients who did not have recurrence tend to have a higher proportion of stromal TILs (p = 0.27, **Supplementary Fig. 12g**). This association was stronger among MSS tumors (p = 0.076) originating from patients without recurrence and this subset of MSS CRCs had 37% more stroma TILS compared to MSS CRCs from patients who did recur (**Fig. 6d)**.

From our study set, we determined if stromal TILs and intra-tumoral TILs had any association with survival. MSI status as a single variable was not associated with survival (**Supplementary Fig. 14a)**. However, the recurrence status significantly impacted survival (p value < 0.001, **Supplementary Fig. 14b**). Additionally, our cohort includes 10 untreated samples, and treatment status did not significantly affect recurrence in this cohort (p = 0.0589, **Supplementary Fig. 14c**).

Next, we categorized the CRCs into high and low stromal TIL groups using Otsu image thresholding (**Methods**). Otsu’s method provides a single intensity level that is used to separate pixels into two classes, foreground and background. We used the method to minimize intra-class intensity variance, or equivalently, by maximizing inter-class variance, applied across all CRC images. This processing provided a mean stromal TIL proportion of approximately 11%.

Survival analysis on these two groups (high and low stromal TIL) revealed that high stromal TIL levels correlated with longer survival (**Supplementary Fig. 14d**), while intra-tumoral TIL levels showed no such association. This stromal TIL-survival relationship was observed among MSS CRCs (p-value=0.038, **Fig. 6e**), with no significant correlation among MSI CRCs (**Supplementary Fig. 14e**). Overall, these findings suggest stromal TIL abundance was a potential prognostic indicator in MSS colorectal cancer.

## DISCUSSION

We conducted a spatial single-cell analysis on stage III colorectal cancer. Our study employed mass cytometry imaging to determine the CRC cell topographic landscape. To our knowledge, this report has the largest number of stage III CRCs ever used for a spatial analysis study. We developed a novel hierarchical spatial analysis framework – it provides three key information patterns extracted from IMC spatial single-cell data, including: 1) proportions of individual cell types; 2) pairwise interactions between adjacent cells; 3) multicellular local tissue architecture in CNs. Then we evaluated how spatial distribution and cellular interactions within the TME and TIME were associated with MSI status and recurrence.

Analyzing single cell spatial data has many advantages in defining specific cell types and spatial organization properties in cancers^45^. This approach provides highly accurate quantitation of different cell types and their spatial organization features in the TME. From our data, we first estimated cell proportion. Then we used the cell proportion in combination with spatial features to characterize the different cell-cell interactions in their local community. Across all CRCs and the162 tissue regions, we identified pairs of cell types that are likely to interact based on their spatial proximity and mediate specific joint functions in the TME.

The metrics of cell proportion and cell-cell interactions were used to determine associations with specific clinical metrics. For example, among patients with recurrence, their CRCs had a high positive correlation between CD8+ and CD4+ T cell abundance. Patients with MSS CRCs and recurrence showed positive correlations with multiple immune cell types, including B cells and T cells and negative correlations between macrophages and cancer cells. In contrast, the analysis of MSI status alone or immune cell proportion alone did not differentiate recurrence status. These results suggest that considering the cell-cell interaction information improved the prediction of recurrence.

We compared the differential enrichment of cell-cell colocalization at single cell resolution across different CRCs, considering variation within an individual tumor as well as those among different tumors. There was higher interaction between cancer cells and CD8+ T cells among MSI-H CRCs compared to MSS CRCs. This result was consistent with previous studies^46,47^ reporting higher lymphocyte infiltration, especially CD3 and CD8+ T cells in MSI-H CRCs. Our study provided more specific and quantitative measures such as the interaction of CD8+ T cells and cancer cells.

We observed greater interactions between cancer cells and proliferative immune cells in MSI-H tumors than MSS tumors. T cells in contact with colorectal tumor cells proliferate at a higher rate compared to those T cells in contact with stromal cells^48^. This phenomenon is related to the increased levels of neoantigens from MSI-H tumors – this leads to stimulation and proliferation of T cells, which we detected as proliferative immune cells. The high level of interaction between CD8+ T cells and cancer cells is an immunologic biomarker of the better prognosis in MSI-H CRCs. Citing a breast cancer example that parallels this result, the interaction between T cells and cancer cells has been associated with upregulation of activation markers^49^. Interactive contact between activated CD8+ T cells and cancer cells has been associated with immunotherapy response in breast cancer patients^49^.

Beyond single cell level interactions, we defined and evaluated features of the local cell community in CRCs. Among CRC patients with recurrence, their tumors showed enrichment of communities with CD8+ T cells and other type of immune cells. The measurement of lymphocyte proportion as a single variable was not informative for determining recurrence compared to TILs in their spatial context. Extending the concept of CNs, we developed a quantitative measurement of intra-tumoral TILs and stromal TILs. Intra-tumoral TILs in MSI-H tumors were significantly higher than what was observed in MSS tumors^15^. For those patients with recurrence, their CRCs had an increase in intra-tumoral and stroma TILs compared to the CRC from patients without recurrence. Also, we found that a greater abundance of stroma tils was associated with prolonged survival in MSS tumors. These results were consistent with the results from another study determining the prognostic value of intra-tumoral TILs and stromal TILs in stage III colon cancers^50^.

We found more TP53+ tumor cells in MSS than in MSI-H samples. This result reflects the significantly higher frequency of *TP53* mutations in MSS tumors (63%) compared to MSI-H tumors (31%)^37^. Mutations in the gene lead to TP53 protein that accumulates in the cells and becomes visible with immunostaining^51^. The cell type proportion comparison suggests that MSI-H samples tend to be more immunogenic than in MSS tissues, for example with more CD8+ T cells, a trend consistent with previous reports and our independent analysis of TCGA stage III COAD cohort. We discovered a rare population of proliferative immune cell population (<1% of total cells), consisting of a mixture signature of CD4+ T cells, T regulatory cells and macrophages. Golby *et al.*^48^ found that a subset of T cells in close proximity or in contact with tumor cells were more proliferative than those in stroma region. In breast cancer, proliferative immune cells have also been associated with better immunotherapy response^49^. Across all samples, the most common immune cells were macrophage and CD8+ T cells. Colon cancer is now recognized as an immunogenic disease^52^ which prompted us to conduct a detailed quantitative analysis of the interaction between immune cells and the surrounding cancer/stroma/immune environment^41^.

To improve the power of our study we acquired tissue regions from the tumor which let us sample more patients with the tradeoff of not being able to capture as much intra-heterogeneity were the entire tumor section sampled. Despite this sampling bias, we accounted for inter-tissue region variability in our statistical testing. For future studies, using a greater number of markers will reveal additional cell types and various functional states.

## MATERIALS AND METHODS

### Samples

This study was approved by the Institutional Review Boards (**IRB**) from Stanford University and Intermountain Healthcare. We analyzed Stage III colon adenocarcinoma tumor samples (n=52) from Intermountain Healthcare (St. George, Utah, USA). Selection criteria and clinical information are described previously^20^. All patients consented to the study and publication of results.

### Imaging mass cytometry analysis

CRCs were prepared from a subset of these collected tumor samples (n=52). Tissue assessment using H&E-stained sections of all 52 tumor samples were performed by a board-certified pathologist and used as a guide for selection of tissue regions during imaging mass cytometry. Tissues were processed for antibody staining. Briefly, slides were baked for 2 hours at 60°C. Simultaneous dewaxing and antigen retrieval was conducted using the Lab Vision PT Module (Fisher Scientific, Hampton, NH) at 96°C in Epredia Dewax and Heat Induced Epitope Retrieval buffer at pH 9 for 35 min. After cooling, slides were blocked in 3% BSA in TBS for 1 h. Samples were incubated with Maxpar® Human Immuno-Oncology IMC™ Antibody Panel Kit (Fluidigm, San Francisco, CA) for 5 h. Samples were then washed 6x with TBS and dried before IMC.

Using IMC Imaging System (Fluidigm, San Francisco, CA)^53^, we simultaneously profiled 16 protein markers for each tissue section, capturing molecular signatures of tissue architecture, cancer cells, and immune cells. All antibodies conjugated with metal were directly procured from Fluidigm (Maxpar® Human Immuno-Oncology IMC™ Antibody Panel Kit). These 16 antibodies were specifically chosen to target cancer, stromal, and immune cell types (see **Supplementary Table 2**). IMC uses laser beam to accurately ablate every 1µm^2^ of tissue region stained with metal-tagged antibodies from the slide samples. The metal tags are then directed to a time-of-flight mass spectrometer for analysis. This technique produces data of exceptional subcellular precision for FFPE tissue sections presented on glass microscopy slides. For each FFPE slide, between 2-8 tissue regions were chosen for in-depth analysis, with an average of approximately 2098 cells per tissue region. The dimensions of these tissue regions spanned between 141 µm x 500 µm and 1121 µm x 1309 µm.

### Data analysis

We first converted the raw IMC data (in the form of .mcd files) to a standard multi-channel image format (OME TIFFs), where each channel signifies the expression of a specific protein staining marker. Using 500-pixel x 500-pixel crops of the original images, we trained a pixel segmentation model in ilastik^54^ to categorize pixels as cell nuclei, cytoplasm, or background. This trained pixel classification model was then applied across the entire full-resolution dataset. We manually reviewed the probability maps for each IMC image and retrained the classifier on the few images where the model was not accurate so that each IMC image was accurately pixel classified into nuclei, cytoplasm or background.

We performed cell segmentation with CellProfiler. Given the variability in the imaging area of IMC, the number of cells extracted from each image fluctuated between 200 and 1600 cells per regions-of-interest. Once cell segmentation was completed, the protein expressions for each cell were determined by averaging the intensity of signals from other channels within the cell boundary, a process conducted using CellProfiler^55^. Cells that were anomalously small (likely due to signal noise) or excessively large (indicating potential overlapping cells) were excluded from further analysis. The comprehensive workflow is depicted in **Supplementary Figure 15**.

### Genomic characterization

We performed whole-exome-sequencing (WES), PCR and IHC to determine the MSI status of tumors from 42 patients (**Supplementary Fig. 2a**). From the WES data we determined MSI status by running MSIsensor^21^ to compute a MSIsensor score and applied an established cut-off of >= 10 to identify MSI high (MSI-H) tumors^56^ (**Supplementary Fig. 2b**). Where multiple assays had been performed for a patient, agreement of MSI status was unanimous.

### Mapping individual cell identities

We performed unsupervised Leiden clustering to identify 22 clusters (**Supplementary Fig. 3a**). This was followed by supervised cluster merging using lineage marker expression profiles (**Supplementary Table 2**) and tissue location to identify 10 cell types.

### Cell type proportion analysis

We used R to compute the proportion of each cell type at the tissue region level and at the patient level. To generate stacked bar plots of the cell type proportions we used the *ComplexHeatmaps* package. We performed statistical testing to compare the proportion of cell types between with different molecular (i.e. MSI) and clinical characteristics (i.e. recurrence).

We leveraged *propeller*^57^ and *limma* in R to rigorously identify statistically significant differences in cell type proportions. We first applied *propeller* to calculate logit transformed variance-stabilized cell type proportions for each region-of-interest. We fitted a mixed effects linear model to the transformed proportions using lmFit in *limma*. We included tumor purity as a covariate as tissue regions that sampled from the same patient may have different cell type proportions and tumor purity depending on the sampled location (e.g., tumor core vs tumor margin) (**Supplementary Fig. 1b, c**). As multiple tissue regions were acquired for each patient, we accounted for replicates from the same individual using the *duplicateCorrelation* function in *limma* and including this a random effect in the linear model. To make comparisons between patients belonging to two conditions, we performed moderated t-tests using *eBayes* in *limma*. p values were false discovery rate (**FDR**) adjusted using Benjamini-Hochberg.

### Cancer genomic validation of cell type proportions

We replicated our cell type proportion testing on a cohort of Stage III colon adenocarcinoma patients from The Cancer Genome Atlas (TCGA). Cell type proportions were collected from previously published CIBERSORTx deconvoluted transcriptomic data^25^. To determine MSI status, we downloaded MSIsensor scores from cBioPortal^58^ and applied the identical cut-off used for our data (**Supplementary Fig. 4a**). To establish recurrence status, we used a standardized dataset of clinical outcomes from TCGA^59^ and considered patients with a disease-free interval event (DFI=1) to have recurred. Ultimately, from the TCGA Stage III colon adenocarcinoma cohort, we selected patients with CIBERSORTx derived cell type proportions and MSI status or recurrence status. We performed same statistical testing as for IMC data, however, as there is only one replicate, we do not need to account for sampling variation with *duplicateCorrelation*.

### Cell type proportion correlations

We computed the cell type proportions at the patient level and calculated the spearman correlation between each pair of cell types with the *rcorr* function from the *Hmisc* package in R. p values were FDR adjusted and only significant correlations with p<=0.05 were plotted as correlograms using the *corrplot* package in R. The same process was used for both IMC and TCGA CIBERSORTx data.

### Cell-cell interaction analysis using a permutation-based approach

After annotating the spatial single-cell data, we sought to identify pairwise interactions between cells neighboring each other. We implemented the permutation-based neighborhood analysis developed for the Histocat software^17^. In this approach, for each tissue region, we first determined pairwise interactions based on proximity between cell membranes identified in cell segmentation. We defined a cell as interacting with a neighboring cell if the distance between the two outer membranes were 6um or less (**Supplementary Fig. 8a**). Next, we counted the number of pairwise interactions between all cell types (**Supplementary Fig. 8b**). We considered the first cell type to the cell type of interest which lies in the vicinity (i.e., neighborhood) of the second cell type. For each pairwise interaction, we divided the number of the interaction by the number of the cell type of interest to calculate the mean interactions for each cell type pair (**Supplementary Fig. 8c**).

To determine which pairs of cell types are preferentially interacting or avoiding each other, we compared the mean interactions we observed against a baseline distribution where the cells are randomly distributed. To estimate this baseline distribution, we randomly shuffled the identity of the cells while maintaining the same no. of each cell type (**Supplementary Fig. 8d**). We repeated this procedure 1000 times and recorded the number of the pairwise interactions for each cell type pair for each random iteration. Finally, the observed pairwise interactions were compared to the baseline distribution through two one-tailed permutations tests (**Supplementary Fig. 8e**). We considered interactions with P < 0.01 to be significant.

### Cell-cell interaction testing using an interaction score

We developed a quantitative interaction score to enable identification of statistically significant interactions between conditions (**Fig 4b**). We applied the same definition of interacting cells as in the Histocat approach. To reducing the noise in the data, we removed tissue regions with fewer than 20 cells of any cell type and focused on 33 interactions involving immune cells. For the interaction score we computed the number of interactions between the cell pair of interest / total number of interactions for each region-of-interest. As the interaction score represents a proportion, we applied the same *limma* framework as described for cell type proportion testing to rigorously identify statistically different cell type – cell type interactions between conditions.

We quantified the center-to-center distance between neighboring cells to inform our definition of cell neighborhoods. To identify neighboring cells, we applied Delaunay triangulation implemented in the *Squidpy*^18^ python package to construct a spatial graph where cells are represented by nodes on the graph and neighboring cells/nodes are connected as edges. We quantified the distances between neighboring cells across all samples. As shown in **Supplementary Figure 17**, the most frequently observed distance between two neighboring cells across all samples as approximately 10 pixels, which is equivalent of 10 μm.

### Identifying cell neighborhoods (CNs)

We defined cell neighborhoods as an index cell and the 10 nearest neighboring cells surrounding it. In very sparse tissues, an index cell’s closest neighbors may still be far away and beyond the ability of cell-cell communication. Thus, we only considered neighboring cells within 40 μm of the index cell to avoid classifying distant cells within sparse tissues are belonging to the same cell neighborhood. The radius threshold was informed by the cell-cell distance analysis. Using Delaunay triangulation to construct a spatial graph of neighboring cells across all samples, we found that two adjacent cells were most frequently 10 μm apart (center to center) and almost all were within 40 μm (**Supplementary Fig. 17**). Accordingly, any cells located outside a 40 radius from a cell’s center were excluded from the count of neighboring cells. Indeed, most cells had ten immediate neighbors.

We expect that CNs with similar cell compositions will have similar functions. Therefore, we performed K-means clustering to identify consistent cell neighborhoods across all patients and found that 8 neighborhoods were the most interpretable yet discriminative. We annotated the neighborhoods based on the composition of cells within the neighborhood.

### Cell neighborhood proportion testing

As each cell has its own neighborhood, we compare cell neighborhood proportions using the same methodology as comparing cell type proportions. Similarly, we compared cell neighborhood proportion correlations using the same approach as for comparing cell type proportion correlations.

### Identification of tumor-infiltrating-lymphocytes (TILs)

We defined two classes of TILs: intratumoral TILs (intra-tumoral TILs) and stromal TILs (stroma tils). We defined intratumoral TILs as lymphocytes (I.e. CD4+ T cell, CD8+ T cell, B cell) that belonged to a tumor cell neighborhood (I.e. bulk tumor CN, TP53+ tumor CN, proliferative tumor CN). Likewise, we defined stromal TILs as belonging to a stromal cell neighborhood (I.e. stromal CN, immune-enriched stromal CN).

With this definition we calculated the proportion of TILs relative to the total number of tumor cells in tumor CNs for each tissue region. To compare between different conditions, we adopt the same framework used for cell type proportion testing.

### Validation of TILs

We downloaded public data of CODEX spatial proteomics^15^ of Stage III and IV colorectal cancers to validate our TILs identification approach. As the cells had already been classified and aggregated into cell neighborhoods, we labelled each T cell or B cell subtype as a lymphocyte and considered those belonging to the CN-2 (Bulk tumor) to be intra-tumoral TILs.

### TILs survival analysis

To determine whether the abundance of TILs was associated with overall survival (OS) we first classified patients as low or high TIL and then applied a cox proportional hazards model. We sought to automatically classify patients are low or high TIL and selected the Otsu threshold method, implemented in the *autothresholdr*^60^ package for R. This approach finds a threshold that maximizes the variance between two classes and minimizes the variance within each class. After identifying low and high TIL patients, we performed survival analysis using the *surviva*l package for R.

## Data availability

The Seurat and Scanpy objects for single-cell data, along with a CSV file containing coordinates and cell types for all single cells, have been deposited in Zenodo under the DOI: https://doi.org/10.5281/zenodo.13901180. The original IMC image data has also been uploaded to Zenodo under the same DOI. All these files are publicly accessible as of the publication date.

## Code availability

All the code used in this study, along with detailed instructions, are available with instructions on GitHub: https://github.com/BiomedicalMachineLearning/CRC_Spatial_Landscape/tree/main

## ACKNOWLEDGEMENT

We would like to thank Ashely Dean for support in the study. We also thank Billy Lau for useful discussion. This work was supported from NIH 5R01CA280089 (HJ.L., H.P.J.), NIH 5P01CA265772 (H.J.L, A.S., X.B., I.W.) and an award from Intermountain Healthcare (HJ.L., H.P.J., A.S., X.B., I.W.). Q.N. was funded by Australia National Health and Medical Research Council (NHMRC Project Grant 2001514), and Australia NHMRC Investigator Grant (GNT2008928). A.S. received support from a Fullbright Scholarship. H.P.J received additional support from the Clayville Foundation.

