## Supplementary Figures for "The single-cell spatial landscape of stage III colorectal cancers"

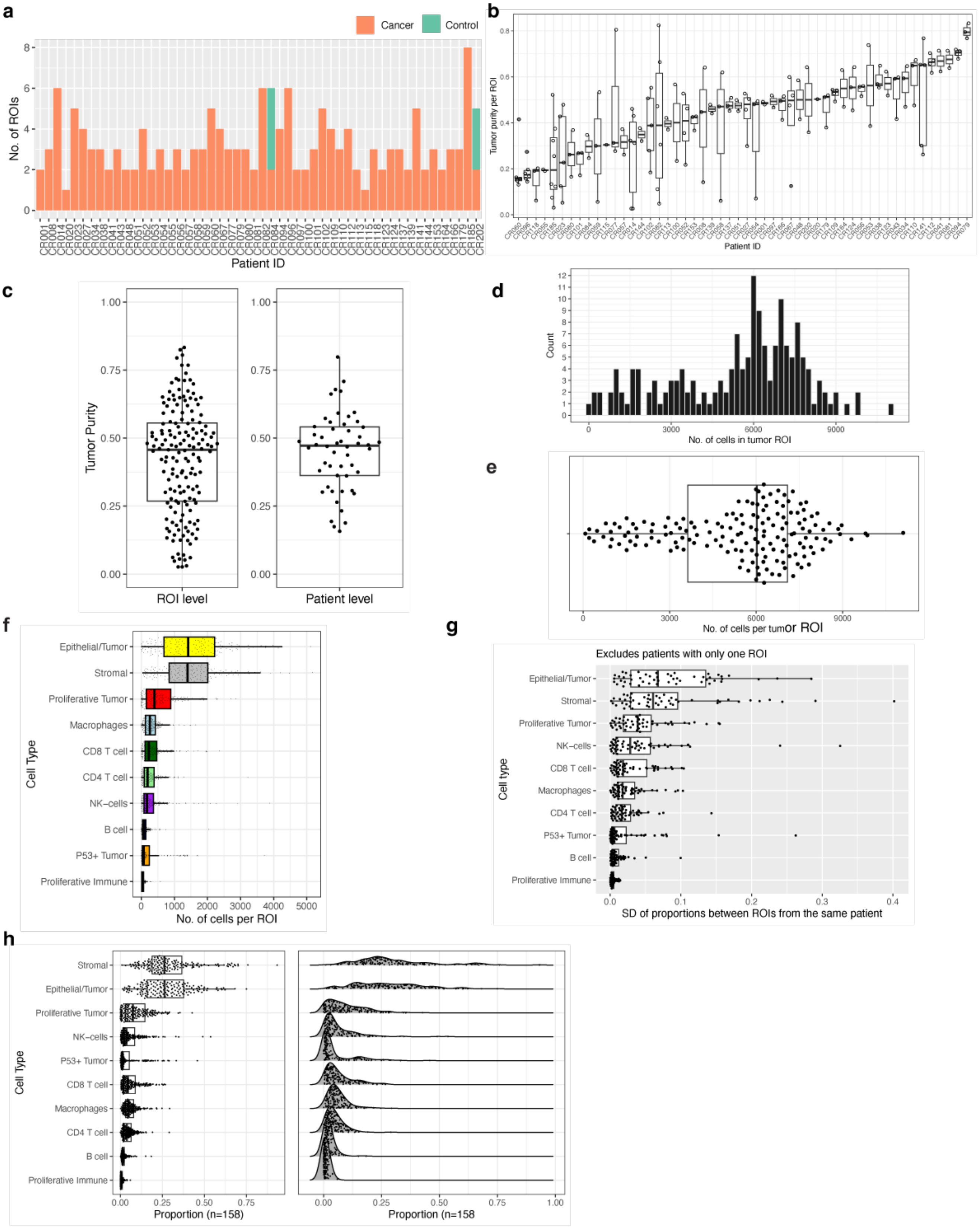


**Supplementary Figure 1**. Dataset description. a) Number of ROIs per sample, b) Tumor purity of ROIs per sample, c)The distribution of tumor purity by ROI and sample, d) Histogram of number of cells across tumor ROIs, e) Boxplot of number of cells across tumor ROIs, f) Number of cells by type per ROI, g) Variability in cell proportion across ROIs within individual sample by standard deviation (SD), h) Boxplot and ridge plot showing the distribution of cell proportion by type across all samples

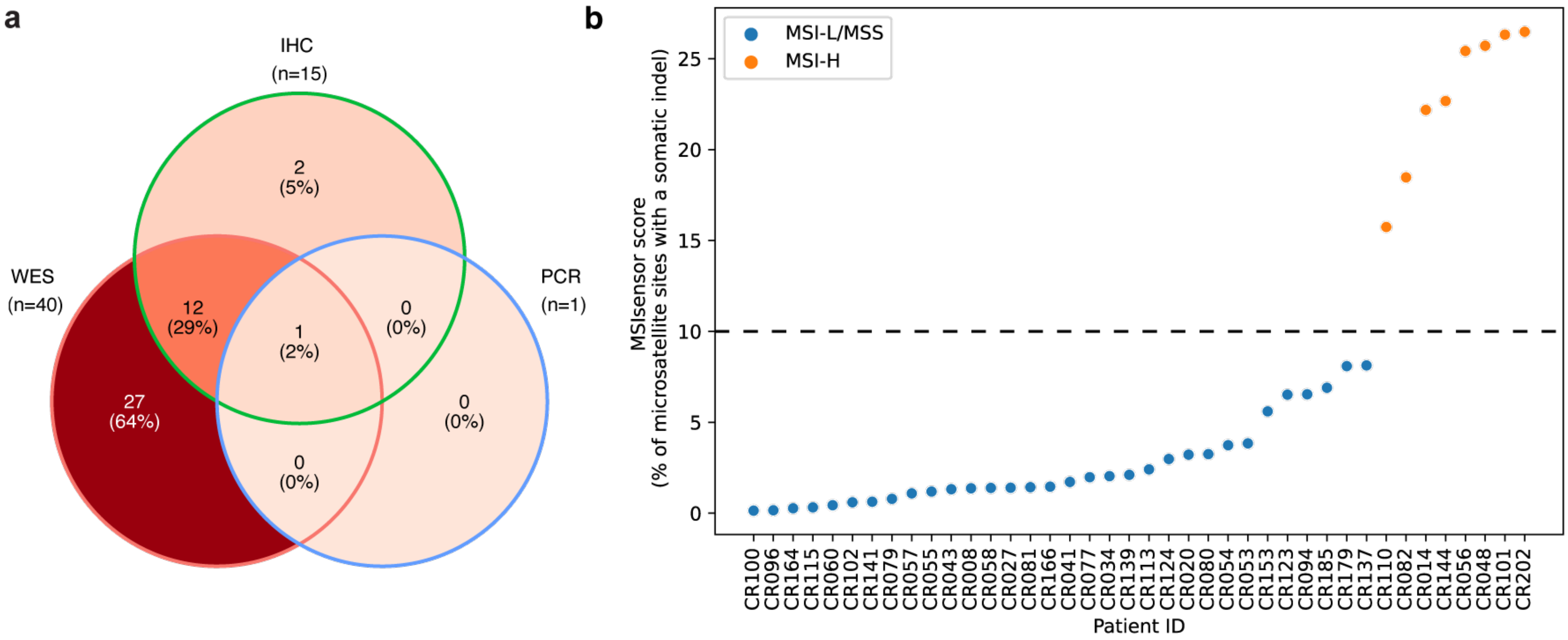


**Supplementary Figure 2.** MSI determination. a) Assays available for determining MSI status. b) MSI status determined by MSIsensor using WES data

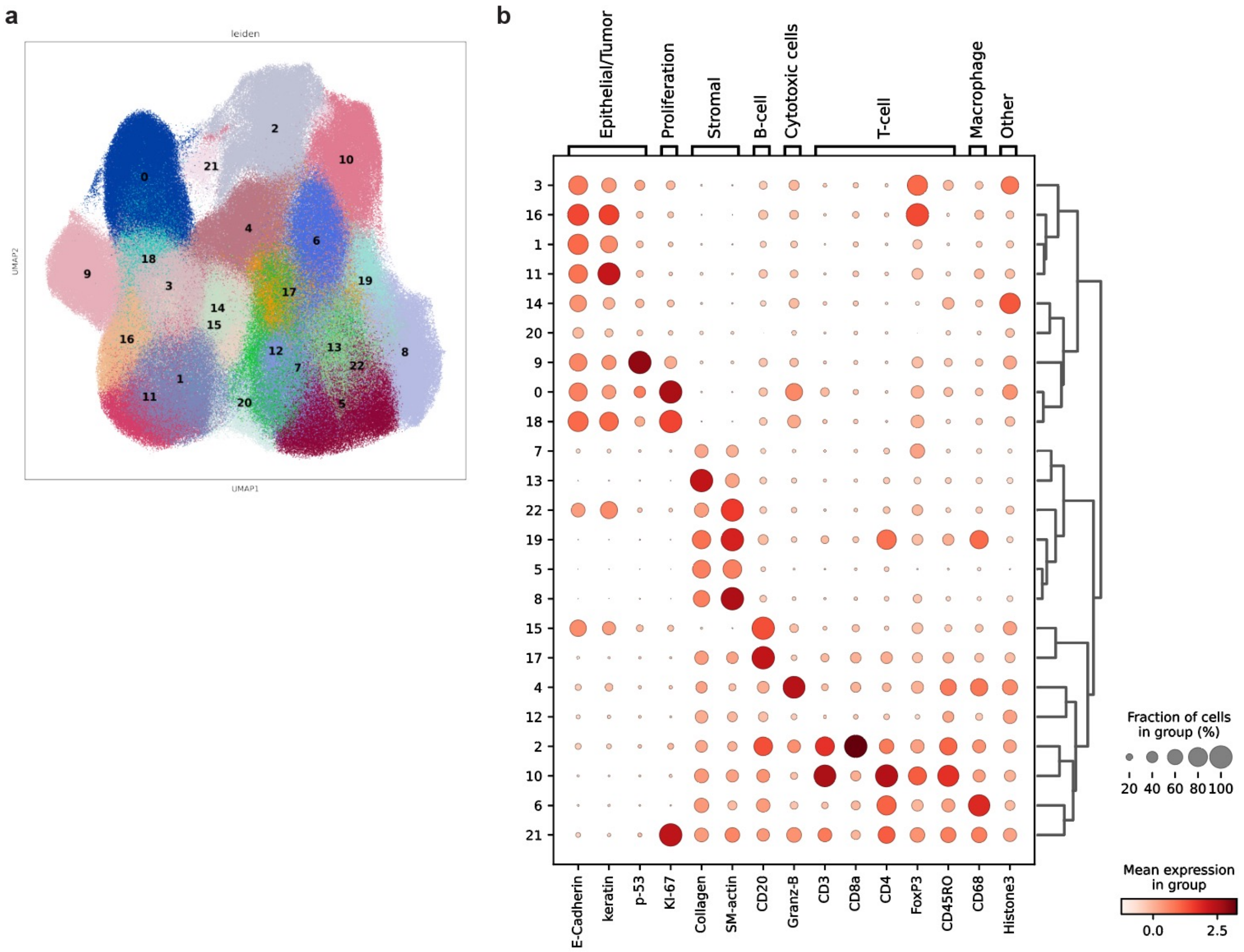


**Supplementary Figure 3.** Cell type clusterings. a) Leiden clustering identified 22 clusters, b) Dot plot of marker expression profile of each cluster

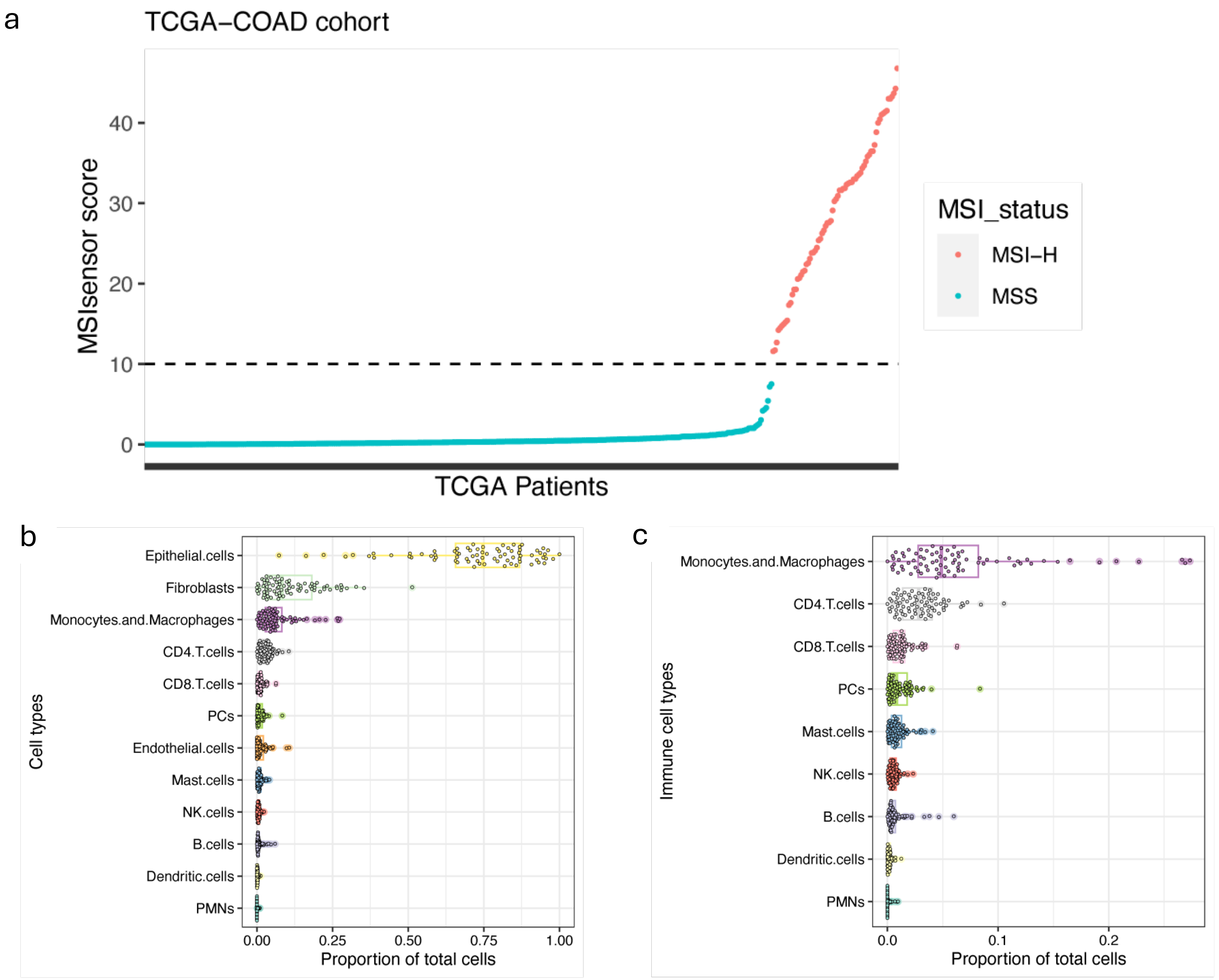


**Supplementary Figure 4.** Cell type proportions across 77 stage III colorectal adenocarcinoma (COAD) patients, as determined by CIBERSORTx deconvolution. a) Microsatellite instability (MSI) status derived from MSIsensor scores based on whole exome sequencing (WES) data, b) Distribution of proportions of all cell types across patient samples, c) Distribution of proportions of immune cell types across patient samples.

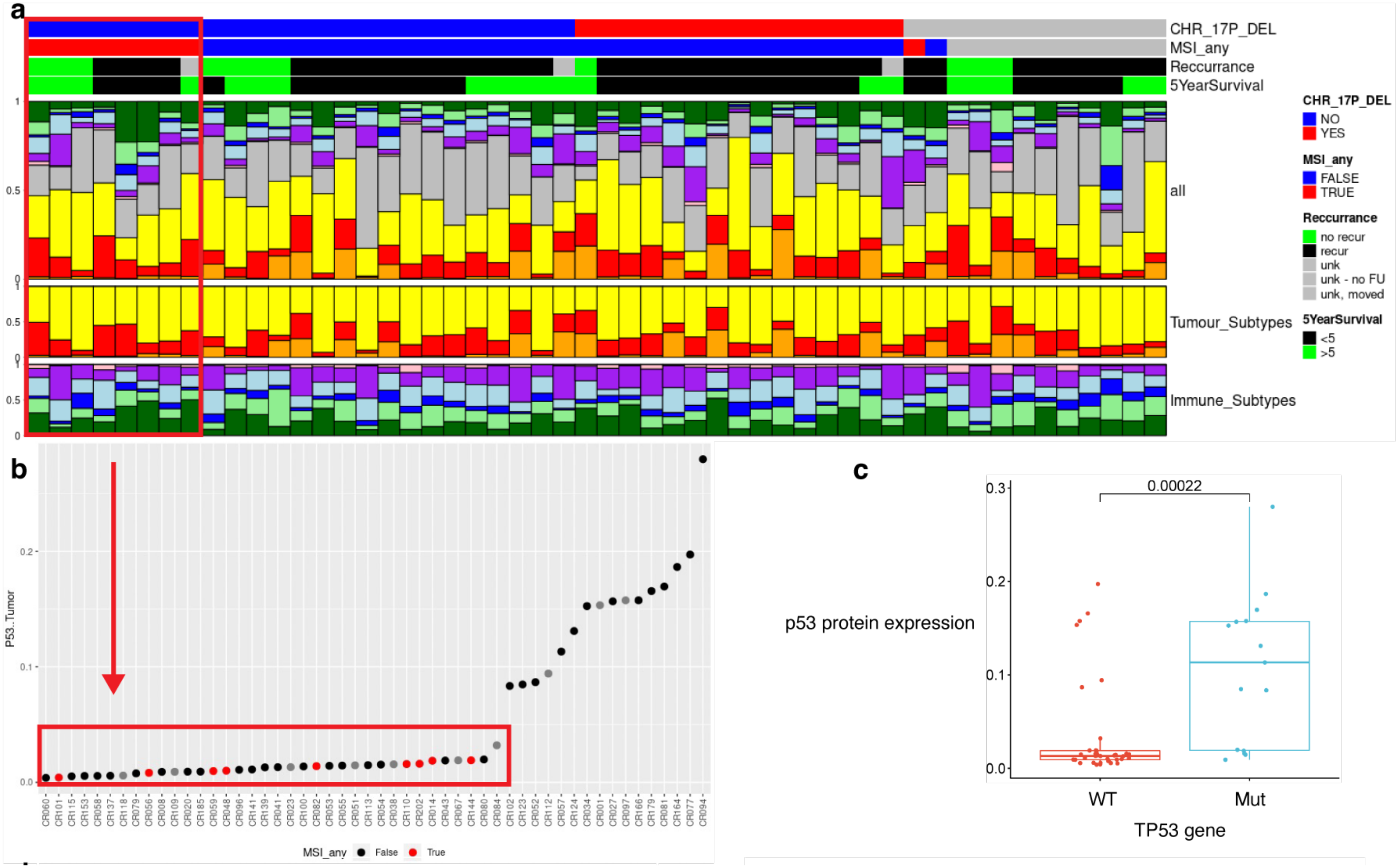


**Supplementary Figure 5.** Comparison of cell type proportions between sample groups based on MSI status and recurrence. (a) Stacked bar plot depicting the cell type composition for each patient, subdivided into tumor and immune cell subtypes, (b) All MSI-H samples exhibited less than 5% p53+ tumor cells, (c) Comparison of p53 protein expression between samples with and without non-synonymous TP53 mutations.

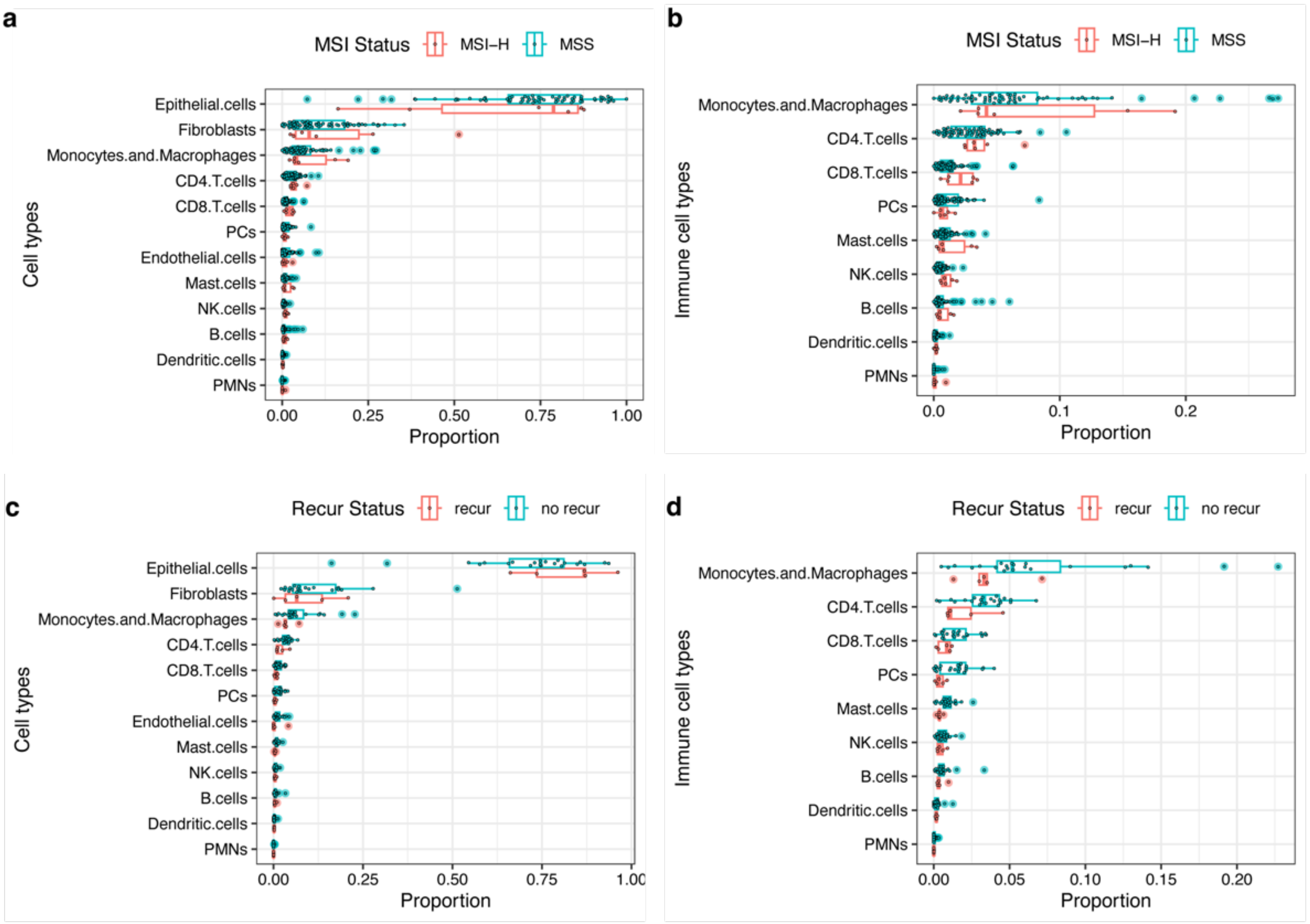


**Supplementary Figure 6.** Comparison of cell type proportions in stage III colorectal adenocarcinoma (COAD) patients from The Cancer Genome Atlas (TCGA). a, b) Comparison of cell type proportions between patients with MSI-H and MSS tumors. c, d) Comparison of cell type proportions between patients who experienced recurrence and those who did not.

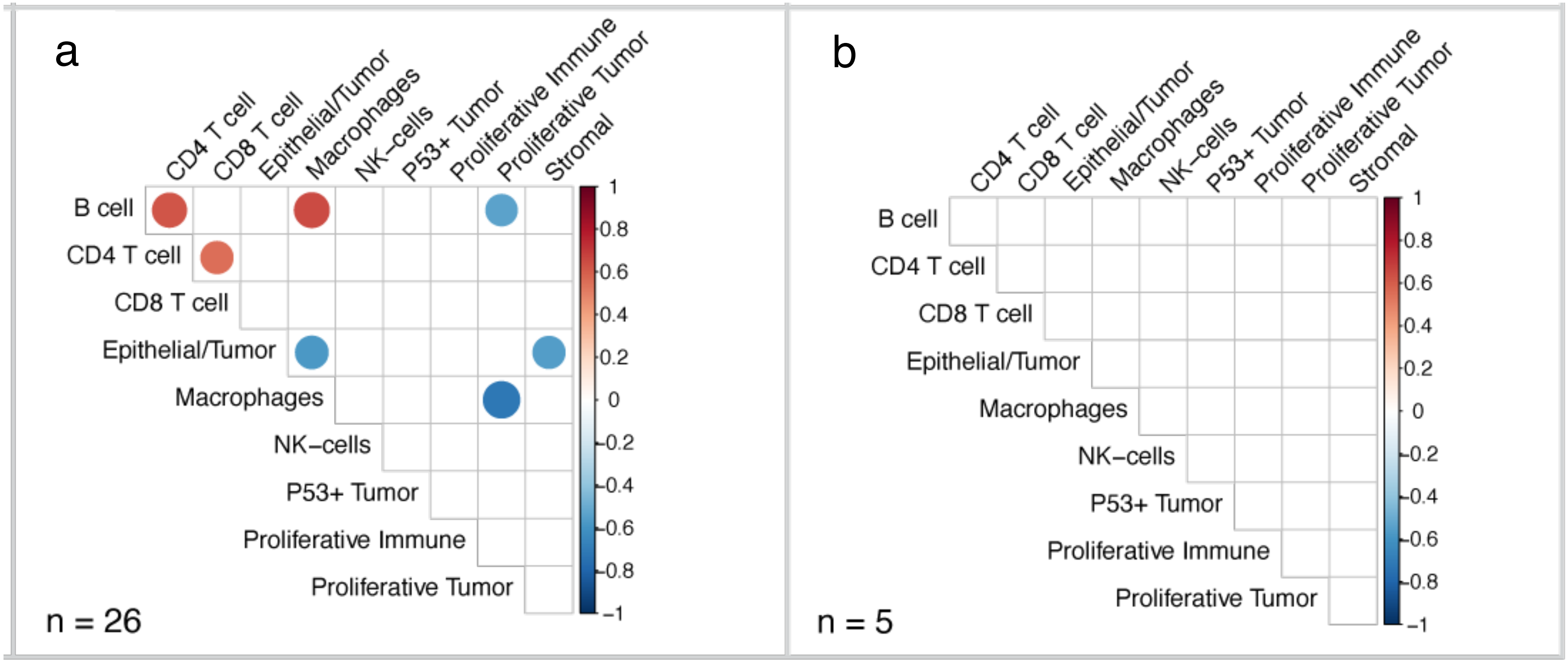


**Supplementary Figure** 7. Significant correlations between the proportions of two distinct cell types in a) MSS recurrent samples and b) MSS non-recurrent samples. All indicated correlations have an adjusted p-value < 0.05.

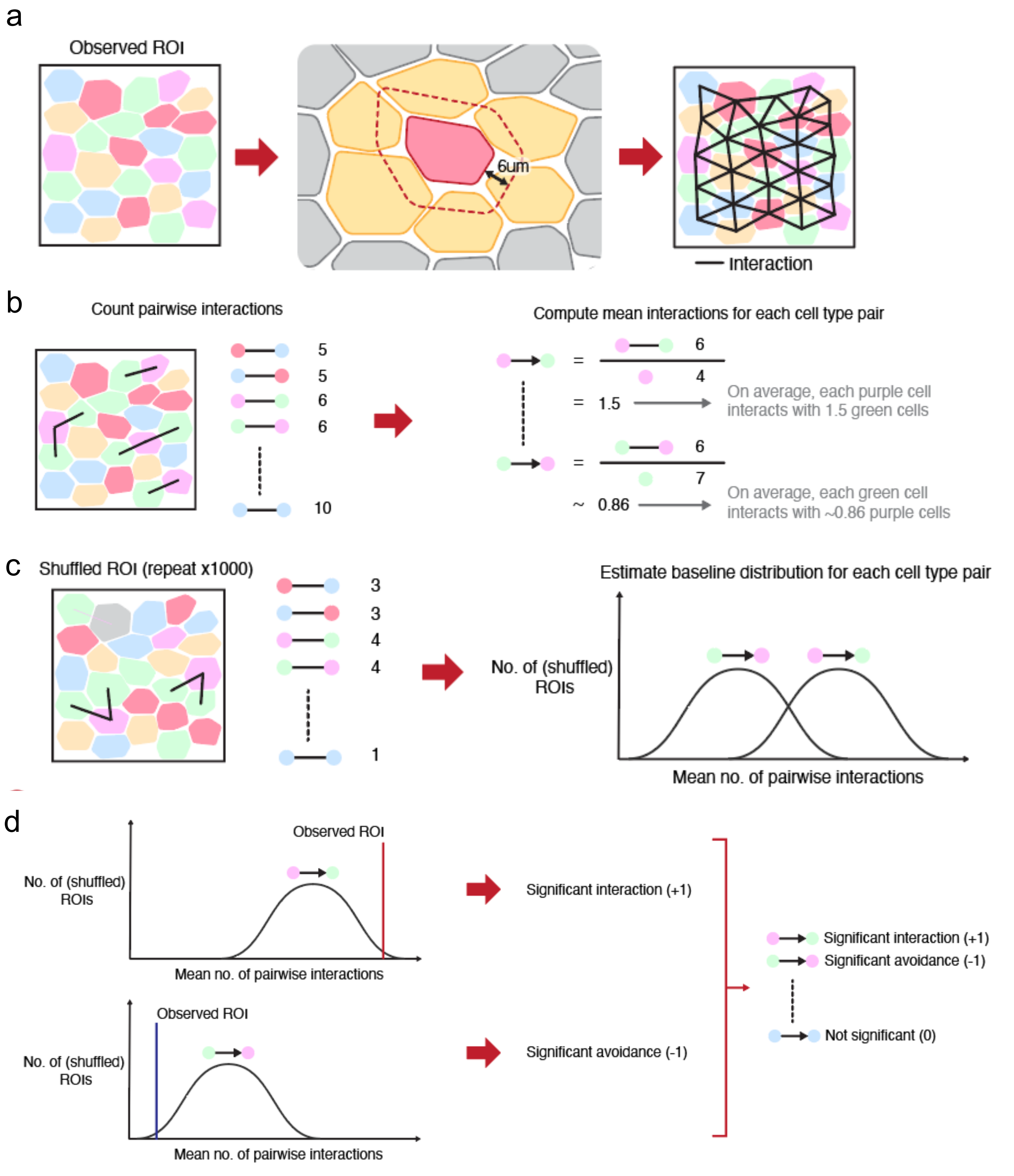


**Supplementary Figure 8.** Overview of the Histocat method. (a) Identification of interacting cells based on spatial proximity, (b) Quantification of pairwise cell-cell interactions, (c) Estimation of the baseline distribution of cell interactions, (d) Statistical significance testing of observed interactions.

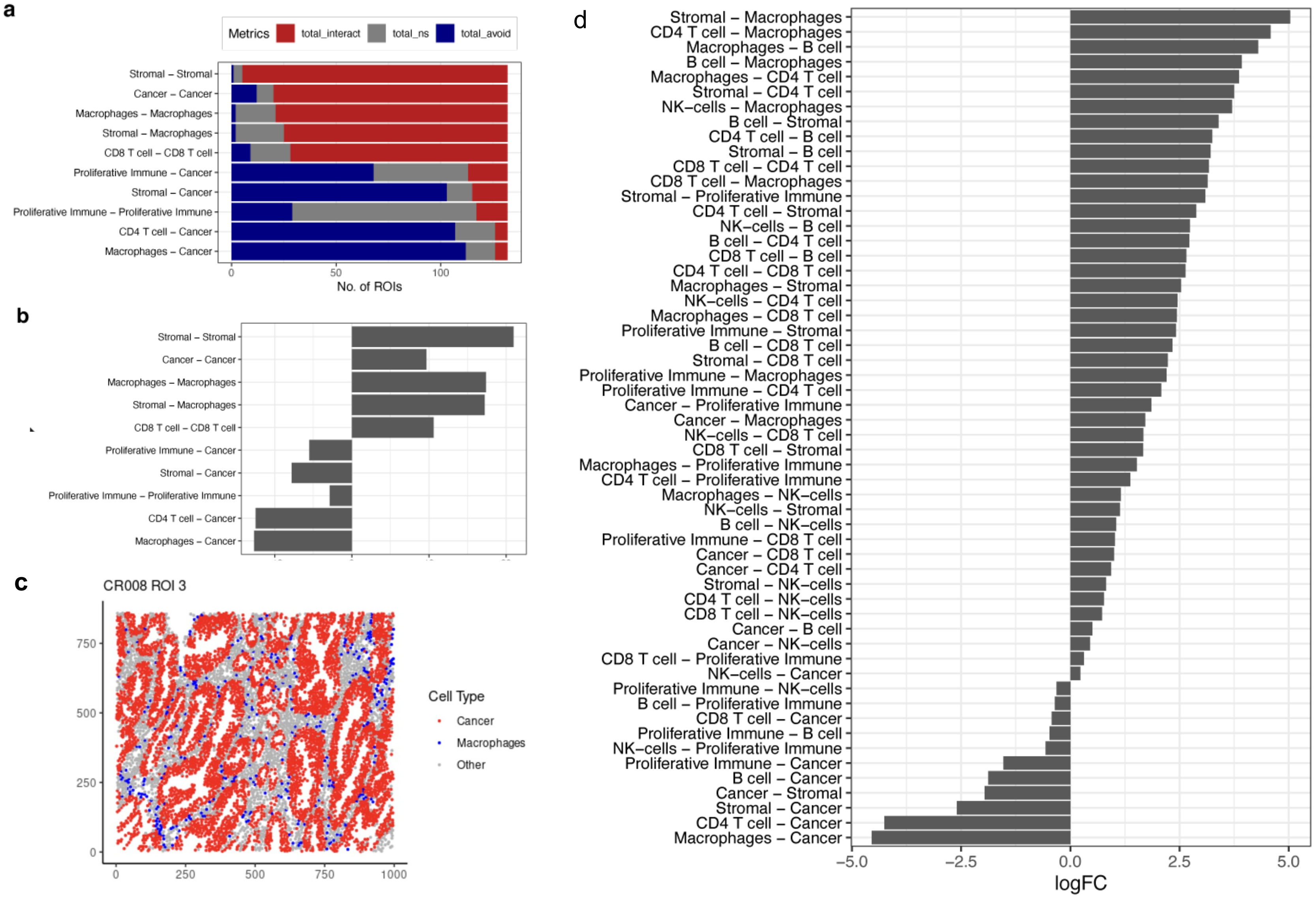


**Supplementary Figure 9.** Significant cell type-to-cell type interactions. a) Number of interacting and avoiding regions of interest (ROIs) for each cell type pair across all tumors, b) Log fold-change (logFC) between the number of homotypic interacting and avoiding ROIs for each cell type pair, c) Example of interaction directionality, illustrating that cancer-to-macrophage interactions differ the most (in terms of logFC) compared to macrophage-to-cancer interactions, d) Log fold-change (logFC) between the number of heterotypic interacting and avoiding ROIs for all possible cell type pairs.

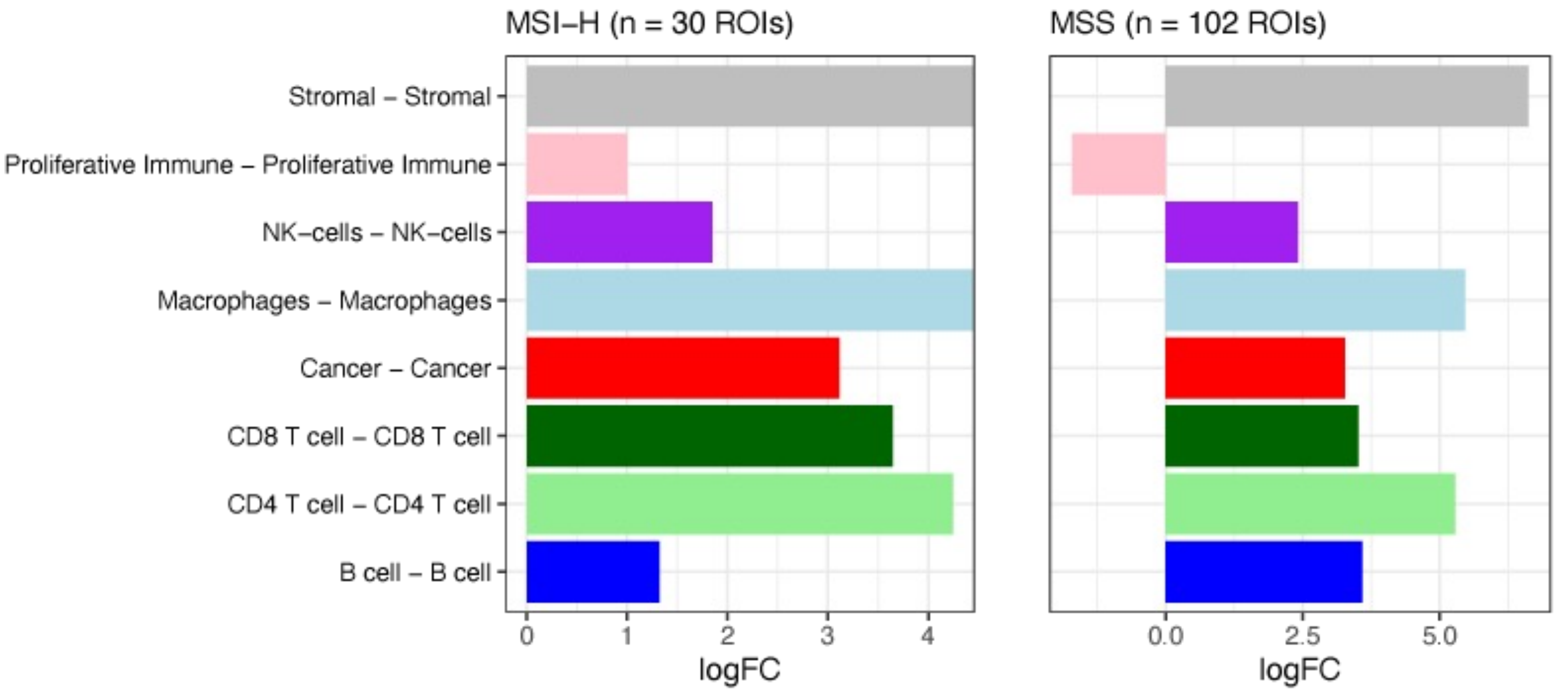


**Supplementary Figure 10.** Homotypic interactions between the same cell types in two groups: 1) MSI-H and 2) MSS. Proliferative immune cell interactions showed opposite directional patterns between the MSI-H and MSS groups.

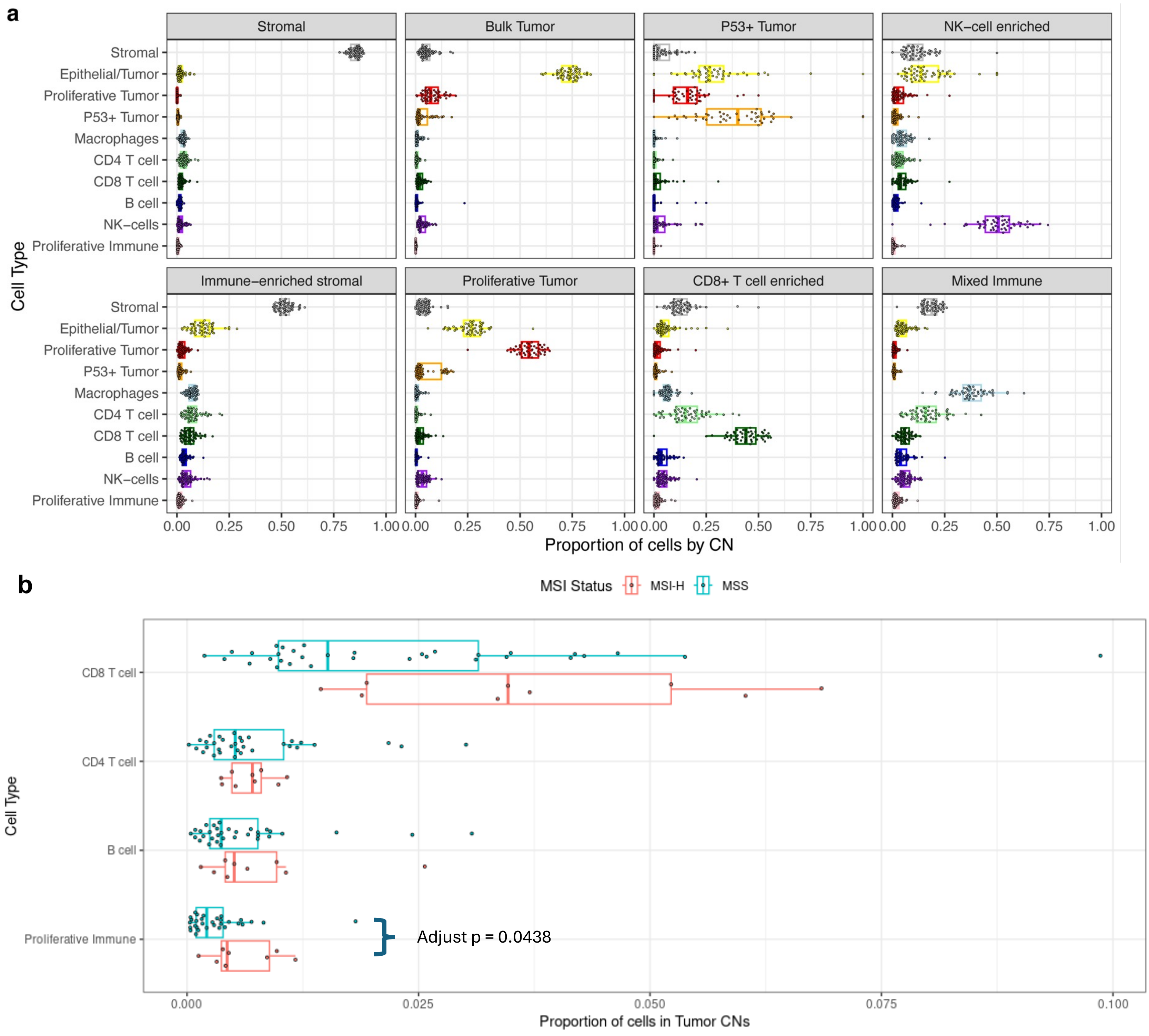


**Supplementary Figure 11**. a) Proportions of different cell types across 8 CNs, (b) differences in cell type proportions within tumor CNs between MSI and MSS samples.

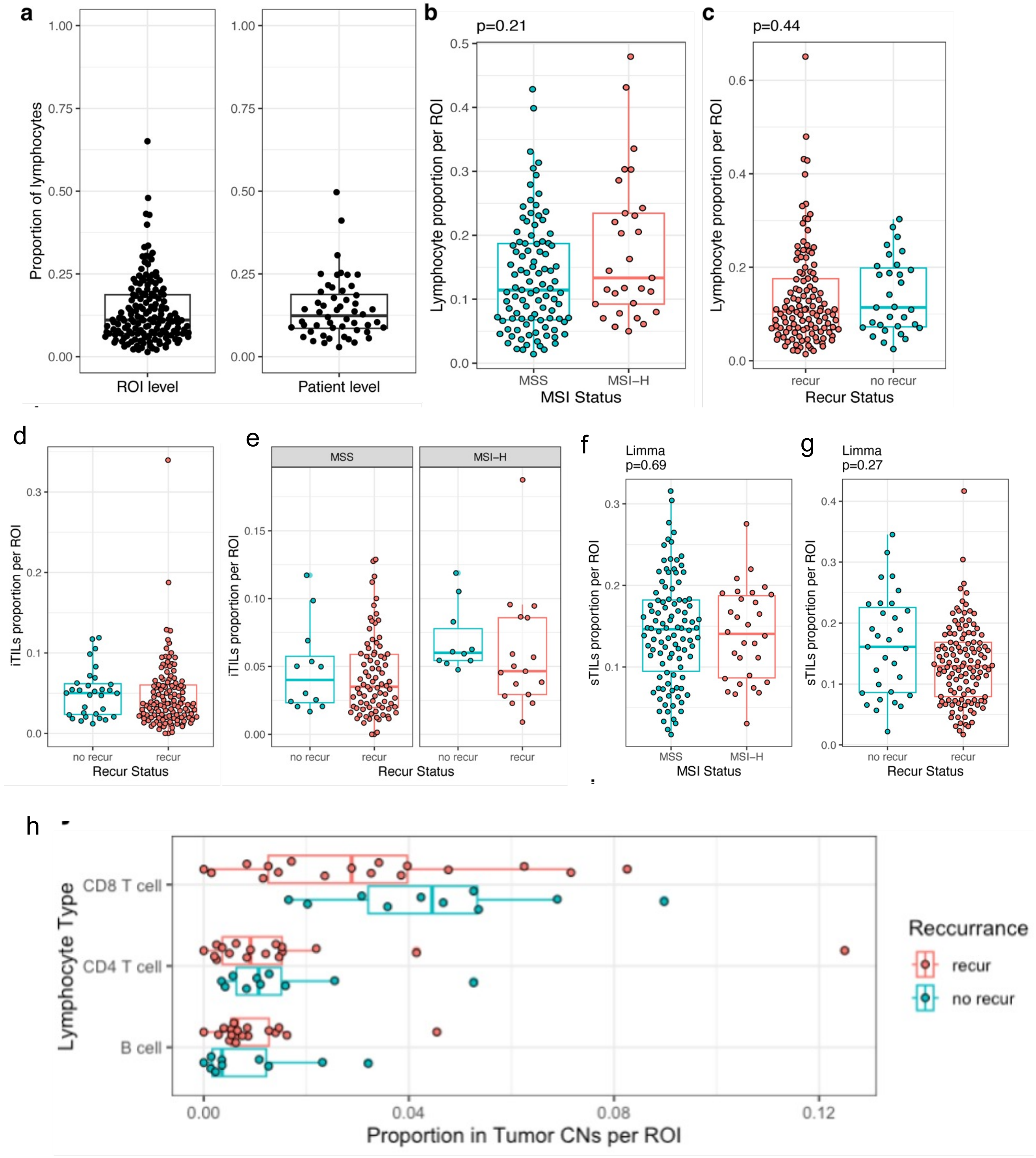


**Supplementary Figure 12.** Comparison of lymphocyte, iTILs, and sTILs proportions. a) Proportion of lymphocytes per region of interest (ROI) across all ROIs and per sample across all patients, b, c) Differences in lymphocyte proportions per ROI between MSI and MSS samples, and between recurrent and non-recurrent samples, d) Proportion of iTILs in patients with and without recurrence, e) Proportion of iTILs in recurrent versus non-recurrent patients, stratified by MSS and MSI-H status, f, g) Differences in sTIL proportions per ROI between MSI-H and MSS samples, and between recurrent and non-recurrent patients, h) Differences in the proportions of three types of immune cells per ROI within tumor CNs between patients with and without recurrence, i) Proportion of iTILs in MSI-H versus MSS patients from CODEX data.

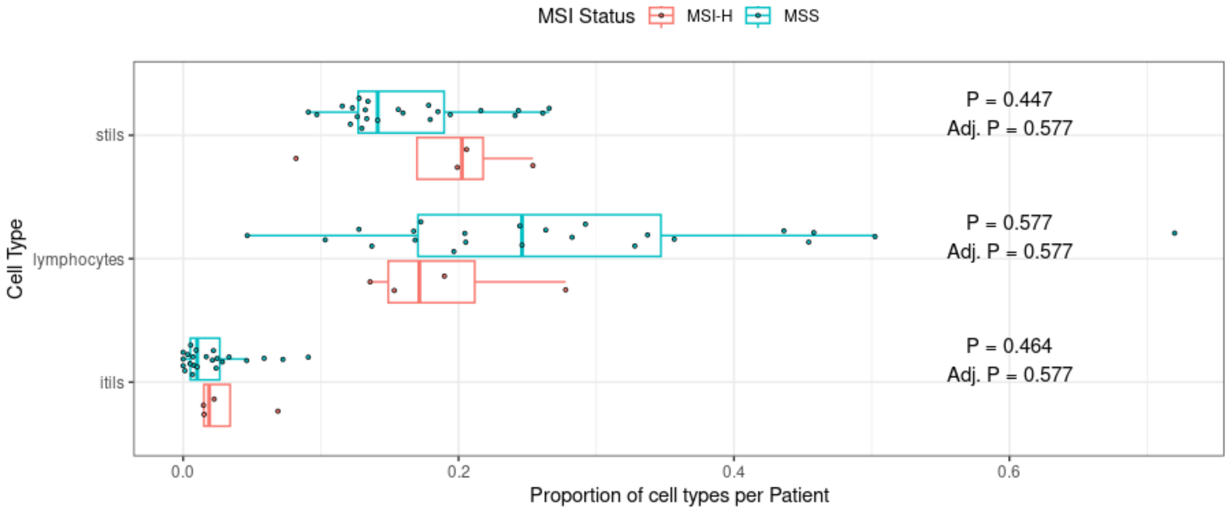


**Supplementary Figure 13.** Comparison of lymphocyte, iTILs, and sTILs proportions from stage III samples (n=27) from CODEX study including 4 MSI-H and 23 MSS CRC: proportion of sTILs (top), lymphocytes (middle), iTILs (bottom) in MSI-H versus MSS patients.

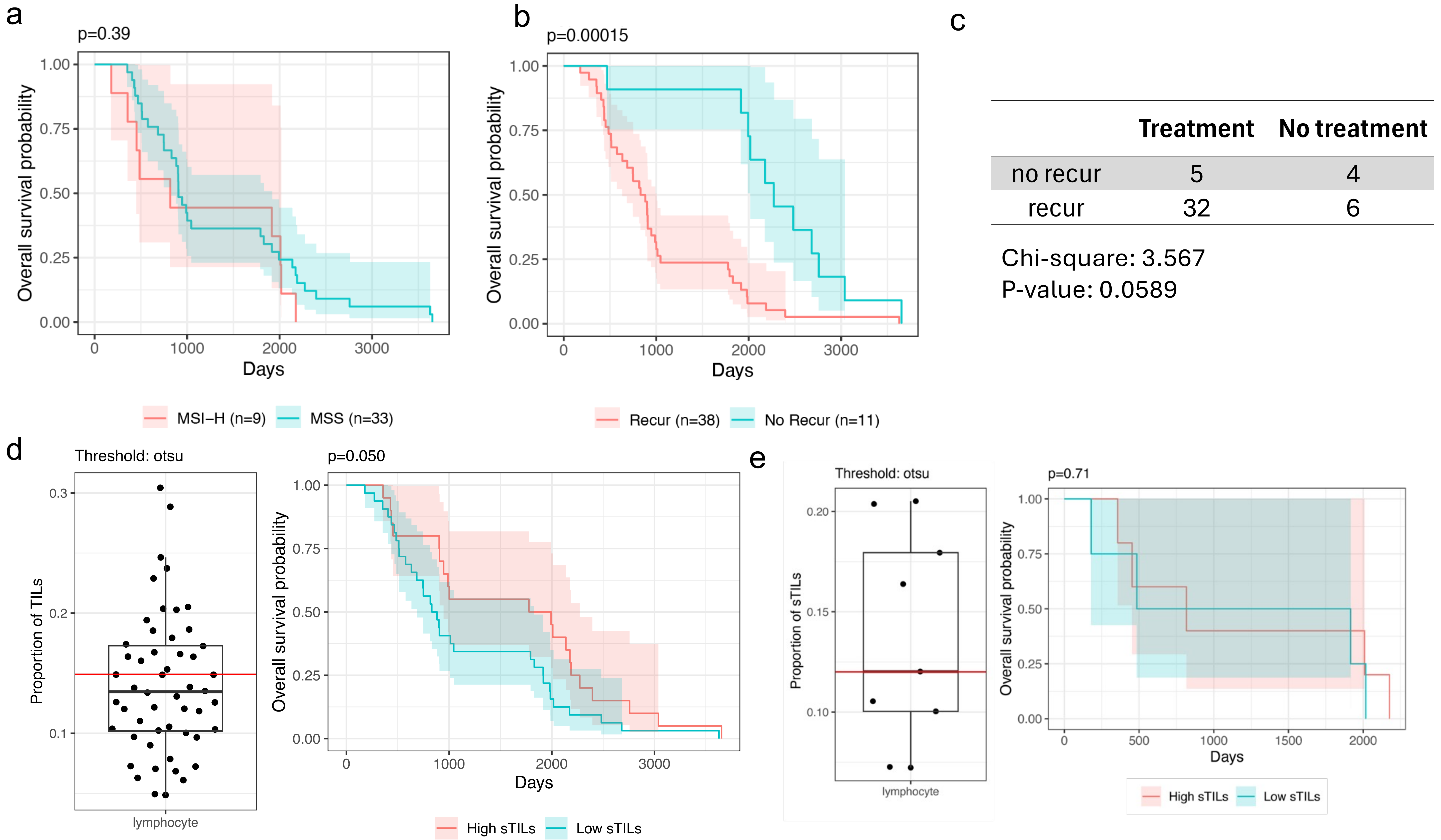


| | Treatment | No treatment |
|---|---|---|
| no recur | 5 | 4 |
| recur | 32 | 6 |

Chi-square: 3.567
P-value: 0.0589

**Supplementary Figure 14**. a) Survival analysis comparing MSI-H and MSS samples, b) Survival analysis comparing patients with and without recurrence, c) chi-square test for treatment vs recur status, d) classification of samples into high and low stroma tils using Otsu thresholding, with corresponding overall survival curves for all samples, and e) only for all MSI samples (n = 9)

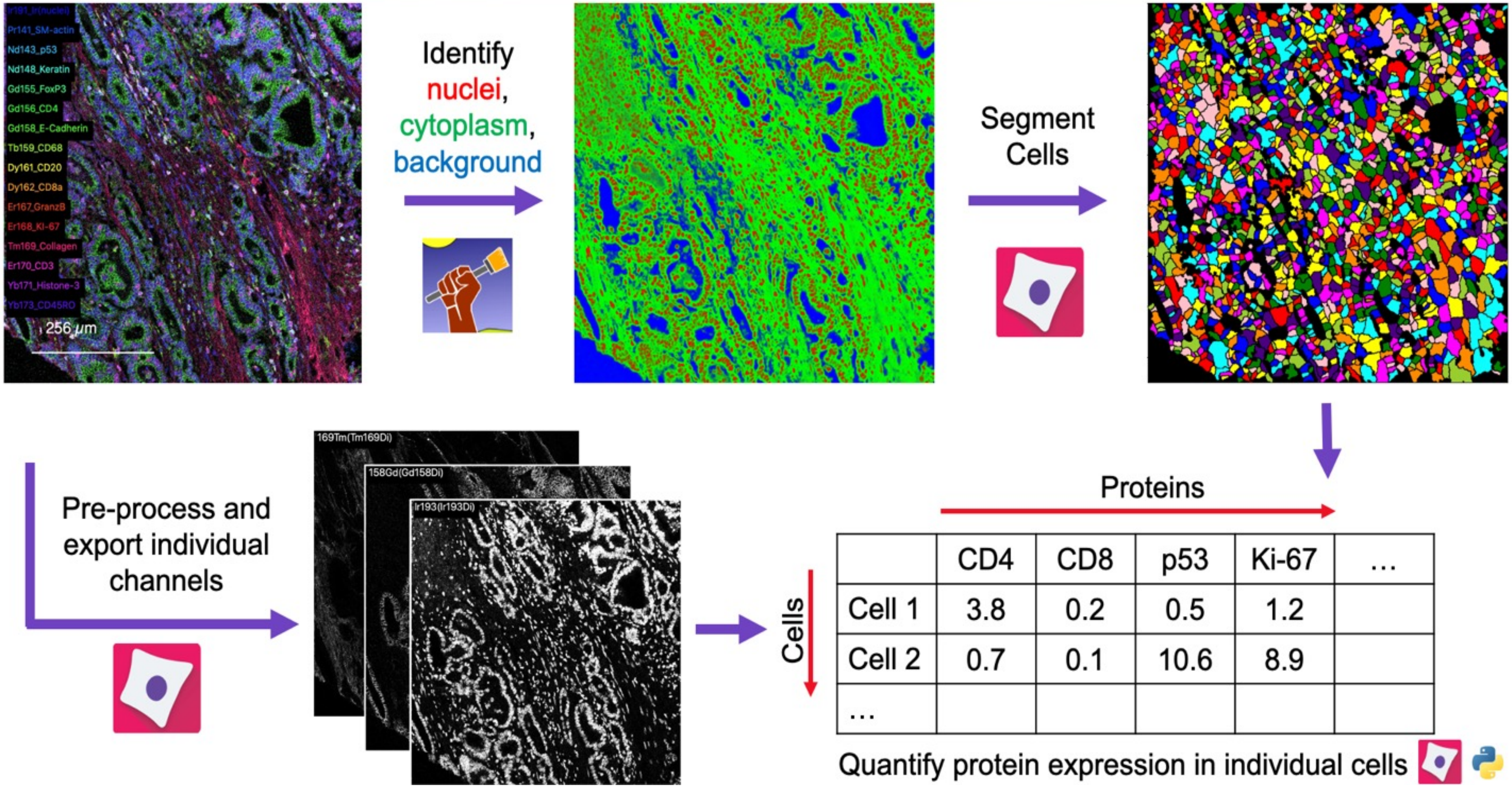


**Supplementary Figure 15**. Overview of cell segmentation and protein expression quantification pipeline

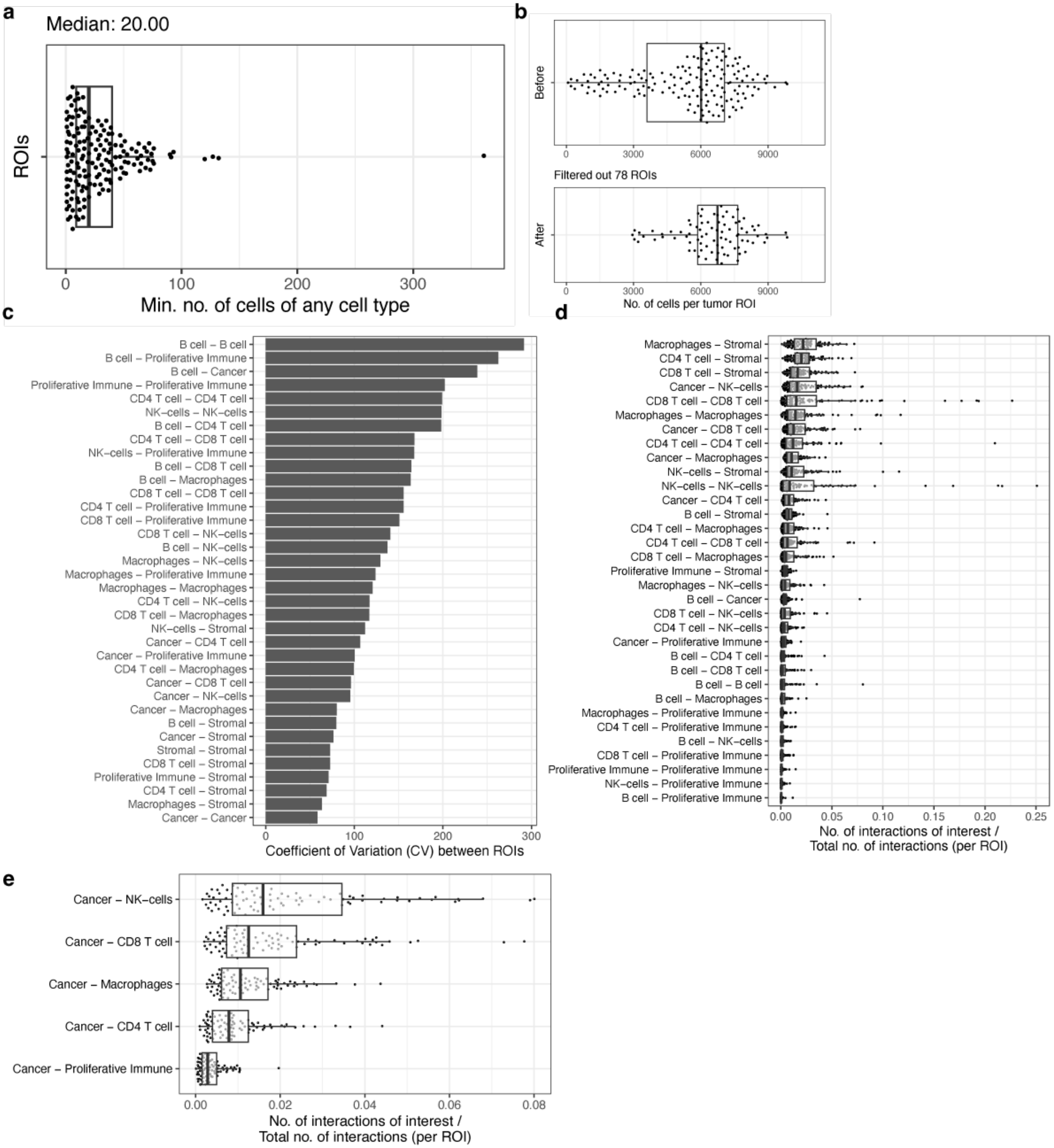


**Supplementary Figure 16**. Interaction score analysis. a) Minimum number of any cell type per region of interest (ROI) across all ROIs, b) Number of cells per ROI before and after filtering out ROIs with fewer than 20 cells of any type, c) Coefficient of variation for pairwise cell type interactions between ROIs, dd) Distribution of the ratio of interactions-of-interest to the total number of interactions in each ROI, (e) Displaying only interactions between cancer cells and immune cell types.

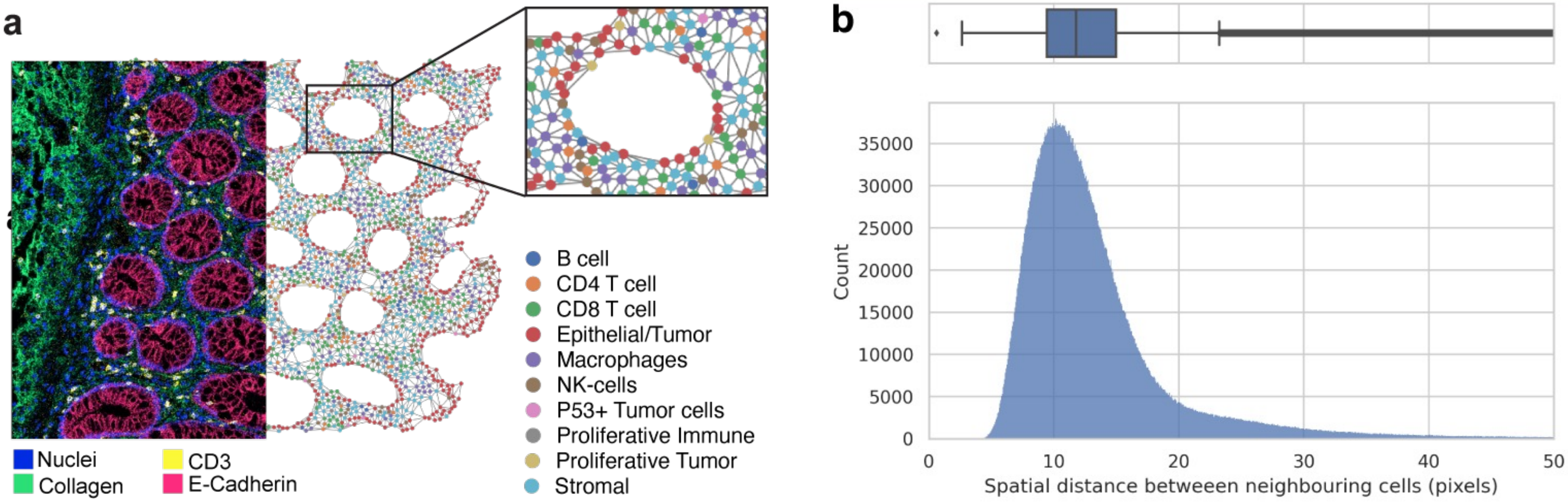


**Supplementary Figure 17**. a) Construction of a spatial graph using Delaunay triangulation, b) Distribution of spatial distances between cells.
